# Advancing global DNA-referencing of bushmeat in African tropical forests: a tool for identifying trade hotspots and dynamics in western and central Africa

**DOI:** 10.1101/2025.05.12.653485

**Authors:** Daniel Pires, Cidalia Gomes, Belinda Groom, Sylvain Dufour, Emmanuel Danquah, Nathalie Van Vliet, Gabriel Ngua Ayecaba, Alain Didier Missoup, Komlan Afiademanyo, Chabi Sylvestre Djagoun, Sery Bi Gonedele, Anne-Lise Chaber, Ayodeji Olayemi, Agostinho Antunes, Philippe Gaubert

**Author notes:** These authors contributed equally to this work (Agostinho Antunes, Philippe Gaubert).

## Abstract

The bushmeat trade in Africa is a largely unregulated activity that drives the unsustainable exploitation of wild terrestrial vertebrates—an issue often coined the bushmeat crisis. We build on a four-gene mitochondrial DNA-typing approach to develop an unprecedented reference database framework for effectively tracing the bushmeat trade in tropical Africa. Our dataset comprises over 2,500 samples collected over a 13-yr period across 10 African countries and two European airports. Relying on an expert analytical pipeline and ∼8,700 nucleotide sequences, we identified 96% of samples to the species-level. In contrast, we estimated that conventional two-gene approaches would have yielded 18–26% erroneous or inconclusive taxonomic assignments. DNA-typing refined > 50% of field identifications, with refinement reaching 92-95% for highly processed carcasses seized in Europe. Leveraging expanded taxonomic representation, we empirically refined the genetic species thresholds applied to bushmeat and provide a reproducible pipeline for using our expert reference database from NCBI. Overall, we identified 133 species—mostly mammals—with one-third listed as of conservation concern, and uncovered cryptic diversity evidence within several taxa. Our results also demonstrated the value of community ecology indexes for large-scale monitoring of the bushmeat trade. While national markets generally mirrored regional biodiversity patterns across tropical Africa, Benin emerged as a notable outlier and a key wildlife trade hotspot. We advocate for the integration of community ecology frameworks into surveillance efforts to guide more effective, regionally tailored mitigation strategies. Continued, standardized sampling is essential to broaden taxonomic coverage and enhance detection of cryptic biodiversity in genetic bushmeat monitoring.

## Introduction

Wildlife trade -the trade of any forms of life sourced from the wild- is a lucrative industry responsible for the yearly movement and selling of millions of organisms across the world, and is a major contributor to the 6^th^ mass extinction (Ceballos et al., 2017; Fukushima et al., 2020; Hinsley et al., 2023). Regulating the wildlife trade has proven a challenging task, notably because of the fuzzy boundary between legal trade and illegal trafficking (Hinsley & Roberts, 2018; ‘t Sas-Rolfes et al., 2019), which itself is greatly influenced by geopolitical shifts (Ribeiro et al., 2022). The wildlife trade encapsulates highly contrasting socio-ecological stakes, from positive impacts on global economies (generating several hundreds of billions US$ / year) and rural household welfare involved in the trade chain, to negative impacts in terms of biodiversity conservation (extinctions) and human health (risks of zoonotic spillovers) (Gallo-Cajiao et al., 2023; Morton et al., 2021; Prasad et al., 2022).

The bushmeat trade in Africa, involving the hunting and sale of wild terrestrial vertebrates from tropical ecosystems, is a marking instance of permissively authorised wildlife trade activities on which rural communities may heavily rely (Friant et al., 2015; Kümpel et al., 2010; Schulte-Herbruggen et al., 2013). The bushmeat trade is valued at approximately US$ 42–205 M per country per year (Davies, 2002; Lescuyer & Nasi, 2016) and occurs through local to global networks reaching the northern hemisphere (Chaber et al., 2023). In the Congo Basin alone, an estimated 4.9 million tons of bushmeat are harvested each year, exerting unsustainable pressure on wildlife populations and causing local extinctions (hence the term “bushmeat crisis”; (Bennett et al., 2007). Additionally, the bushmeat trade has significant public health implications, as it is a known vector for zoonotic diseases, including Ebola and HIV (Leroy et al., 2004; Peeters et al., 2002), underscoring the urgent need to understand the dynamics of the trade and apply sustainable management practices.

The bushmeat trade in Africa remains a largely unregulated global activity, encompassing the hunting and sale of approximately 500 species of terrestrial vertebrates, predominantly mammals, but also including birds and reptiles (Redmond I. et al., 2006). The overexploitation of bushmeat species, in conjunction with widespread habitat destruction, has already resulted in local extinctions and population declines, particularly among large herbivores, primates, and pangolins (Ingram et al., 2017; Walsh et al., 2003; Zanvo et al., 2020). Given that the bushmeat trade affects a broad range of functional groups, such as seed dispersers, prey regulators, pollinators, and browsers (Djagoun et al., 2023), it is expected to trigger cascading ecological effects within tropical ecosystems (Ripple et al., 2016). This is especially concerning in regions identified as trade ‘hotspots,’ where bushmeat markets are concentrated collapse is likely, driven by the removal of key species with critical ecological functions (Tagg et al., 2019). Despite the significant conservation implications, surveys assessing the sustainability of the bushmeat trade in tropical Africa have been hampered by methodological limitations, including insufficient protocol design, spatio-temporal limitations, and the lack of accurate species identification (Groom et al., 2023; Morton et al., 2021).

The use of DNA-based techniques (or DNA-typing) has revolutionised bushmeat trade surveys by enabling accurate species identification (Alacs et al., 2010), even from processed or smoked carcasses where morphological identification is difficult (Minhós et al., 2013; Olayemi et al., 2011). Furthermore, DNA-typing can provide insights into the geographical origin of bushmeat, helping to pinpoint poaching hotspots and track trade routes (Ghobrial et al., 2010; Zanvo et al., 2022). Most bushmeat surveys have traditionally relied on a single-gene barcoding approach (Minhós et al., 2013; Schilling et al., 2020), focusing on the universal barcode region of Cytochrome Oxidase I (COI) or on the Cytochrome b (Cytb), mitochondrial DNA (mtDNA) genes well-represented in vertebrate reference databases. However, the single-gene barcoding approach has known limitations (Ruedi et al., 2023; van Velzen et al., 2012), which may be exacerbated by additional biases —such as false positives and poor PCR efficiency / sequencing quality— due to factors specific to the bushmeat context, including cross-contamination at market stalls and DNA degradation from dead or processed carcasses. COI, in particular, has been shown to be prone to significant levels of PCR bias in bushmeat species (Gaubert et al., 2015), compounded by reference databases that are often less taxonomically comprehensive than those for Cytb (Ruedi et al., 2023).

While some recent studies have adopted a two-gene approach for DNA-typing in bushmeat trade surveys (Deagle et al., 2014; Félix et al., 2021), we argue that this effort is still insufficient, particularly because conflicting results between the two genes will prevent the taxonomic identification of carcasses. To address these challenges, we have implemented a four-gene DNA-typing approach in our bushmeat surveys for nearly a decade (e.g., Chaber et al., 2023; Din Dipita et al., 2022; Djagoun et al., 2023; Gaubert et al., 2015; Gossé et al., 2022). This method provides a more comprehensive framework and sufficient genetic information to robustly assign species identities to bushmeat samples and unravel cryptic diversity. The primary objective of our study is to implement a global DNA reference database framework that promotes the accurate identification of species and geographic lineages trafficked in the bushmeat trade across tropical Africa. The first specific objective is to empirically evaluate the performance of the four-gene DNA-typing approach, leveraging an unprecedented sampling effort across western and central Africa, including comparisons with single- and two-gene approaches, as well as morphological identifications, while incorporating reassessed gene-specific species delimitation thresholds. The second objective is to assess the effectiveness of the DNA-typing approach as a surveillance tool for monitoring bushmeat trade dynamics across tropical Africa, applying community ecology principles to identify trade hotspots.

## Material and Methods

### Data context

The dataset includes a total of 2,516 genetic samples collected over a 13-year period (2008-2020) across 10 tropical African countries and from two seizures at European airports, as part of a long-term collaborative effort led by PG and collaborators (Chaber et al., 2023; Din Dipita et al., 2022; Gossé et al., 2022; Olayemi et al., 2011) (Fig. 1a). The sampling strategy in Africa opportunistically relied on bushmeat market vendors and hunters, who were informed of the objectives of the study before allowing genetic sampling. Actors were not given incentives or remuneration for species of interest. In European airports (Zaventem in Brussels, Belgium, and Roissy in Paris, France), seizures were conducted as part of joint operations involving national customs and police services. The dataset spans across 19 vertebrate taxonomic orders and 171 putative species or higher-level taxa representative of the bushmeat trade, including mostly mammals (*N* = 153) but also birds (*N* = 6) and reptiles (*N* = 11) (Fig. 1b). Primary taxonomic hypotheses were generated after a variety of protocols and support, depending on survey sites and period (Kingdon 1997 was the major reference).

**Figure 1.**
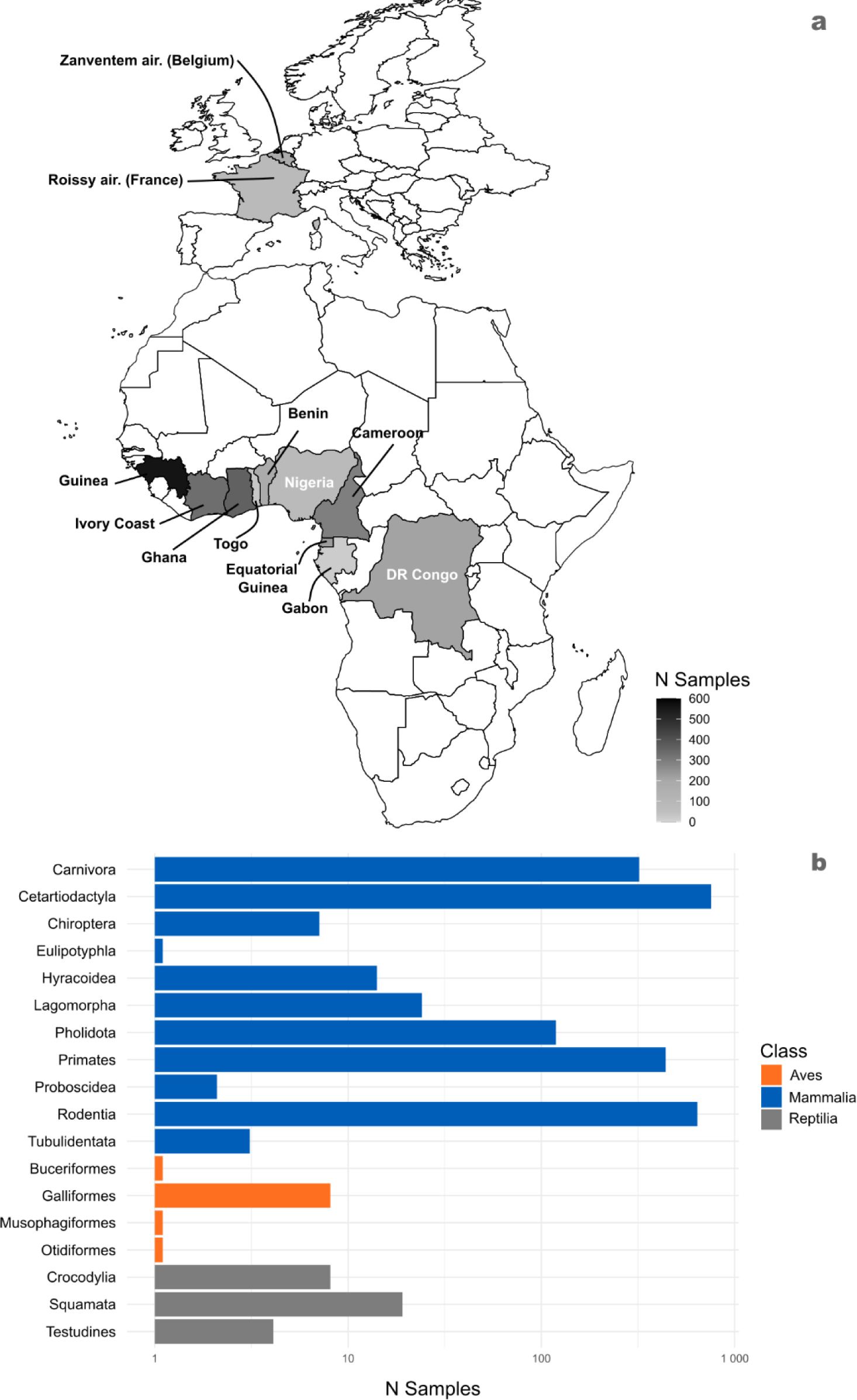
Distribution of the bushmeat sampling effort across countries (a) and taxa (b). Maps were created with package *rnaturalearth* in RStudio.

The samples were DNA extracted, PCR-amplified and Sanger-sequenced for four mtDNA gene fragments following the DNA-typing protocol of (Gaubert et al., 2015). The four markers encompass Cytochrome c oxidase I (COI; 658 bp), Cytochrome b (Cytb; 402 bp), and 12S (167 - 404 bp) and 16S (210 - 528 bp) ribosomal DNA. The original dataset produced as part of this study represents 1,253 genetic samples and 4,422 nucleotide sequences (Genbank accession numbers xxx-xxx), collected from six countries and one European airport (see Table S1, Supplementary information). The remaining data was obtained from previously published works (Chaber et al., 2023; Din Dipita et al., 2022; Gossé et al., 2022; Olayemi et al., 2011), representing 1,263 samples and 4,270 nucleotide sequences.

### Sequence quality and assignment procedure

Sequence quality and taxonomic assignment procedure were conducted on the newly produced dataset of sequences (N=4422). For COI and Cytb, we checked for the presence of potential pseudogenes (or NUMTS; Lopez et al., 1994), by screening for the disruption of their reading frame in Geneious prime v. 2023.3.1 (Kearse et al., 2012). For 12S and 16S, due to the absence of a reading frame, potential pseudogenes were screened for through long-branch and aberrant phylogenetic branching patterns (Triant & DeWoody, 2007). The sequences representative of 12S and 16S were first aligned with Muscle (Edgar, 2004) in Mega v.11 (Tamura et al., 2021) under default settings. Second, we used Gblock (phylogeny.lirmm.fr/phylo_cgi/one_task.cgi?task_type=gblocks), with both relaxed and default settings depending on the levels of divergence within orders, to remove poorly aligned positions, resulting in final alignments of 288-331 bp for 12S and 322-485 bp for 16S. Neighbour Joining (NJ) trees (Saitou & Nei, 1987) were generated with MEGA under the K2P (Kimura, 1980) model and the option “pairwise deletion”. Node robustness was assessed through 500 bootstrap (Felsenstein, 1985) replicates. The same NJ tree approach was used for COI and Cytb as a secondary assessment of sequence quality, as pseudogenes do not necessarily translate into reading frame disruption.

The taxonomic assignment of the newly produced sequences followed a two-step process. First, the sequences were blasted in DNAbushmeat, an expert-curated local database referencing 60 bushmeat species (Gaubert et al., 2015), and were sorted according to their similarity values being equal / superior or inferior to the thresholds empirically established by the developers of the database (COI-Cytb: 95%; 12S-16S: 97%). The taxonomic assignment of the sequence was determined according to the range of species provided by the database, for example, if the best hit was for species A from 99% to 94% and no other species was found within that interval, species A would be the sole assignment. However, in cases where there was another species (species B) within the range of species A, then the taxonomic assignment would be “species A/B”, provided that species B had a best hit ≥ 95-97%, depending on the gene targeted. Second, whenever (i) best hit similarity values were less than the thresholds and/or (ii) a final consensus was not reached among the four-gene taxonomic assignments, a blast search was conducted on the NCBI Nucleotide Blast web server (https://blast.ncbi.nlm.nih.gov/Blast.cgi) using megablast (Sayers et al., 2022). Whenever best hit similarity values were higher than those obtained from DNAbushmeat, taxonomic assignments and final majority consensus were updated. Final ID was reached by checking for the occurrence of the species in the country of interest on the IUCN Red List of Threatened Species (https://www.iucnredlist.org/). The IUCN Red List was also used to extract the conservation status of the identified species. Whenever the taxon in question had a complex taxonomy (e.g., Cercopithecidae), the Mammals of Africa (Kingdon et al., 2013) was utilised to double-check taxonomy and confirm geographic distributions.

In a few cases where taxonomic assignment did not match geographical distribution, final ID followed the original morphological-based field ID (1.5 % of the samples). Final IDs were made to the species or genus level (in case of multiple-species assignment with different species occurring in the same country), and were scored “NA” whenever assignments proved inconclusive (see Table 1). Final validation of taxonomic ID was reached through the assessment of sequence clustering on gene- and order-specific Neighbor Joining trees ((Saitou & Nei, 1987); see Figs S1-S27, supplementary material), which were reconstructed following the above-mentioned settings.

**Table 1.** Multiple-scenario decision table for the final taxonomic identification of bushmeat samples.

| Field ID | COI | Cytb | 16S | 12S | Molecular<br>consensus | Occurrence<br>in country | Final<br>ID | Note |
| --- | --- | --- | --- | --- | --- | --- | --- | --- |
| A | A | A | A | A | A | A | A |  |
| A | A | B | A | A | A | A | A |  |
| A | A | B | C | D | A/ B/ C/ D | At least two of<br>the species | A | Field ID<br>validates<br>species A |
| A | A/B | A | B | A/B | A/B | At least two of<br>the species | Genus<br>of<br>species<br>A and<br>B | Both species<br>belong to<br>the same<br>genus |
| NA | A | B | C | D | A/ B/ C/ D | A | A | Geographica<br>l distribution<br>validates<br>species A |
| Genus X | A | B | C | D | A/ B/ C/ D | At least two of<br>the species<br>from genus X | Genus<br>X | All the<br>species<br>belong to<br>genus X |
| NA | A | B | C | D | A/ B/ C/ D | At least two of<br>the species | NA | Several<br>genera<br>involved. No<br>final ID |

A - Species A
B - Species B
C - Species C
D - Species D

### Assessing the accuracy of DNA-typing taxonomic assignment and morphological-based identification of bushmeat species

Accuracy of the four-gene DNA-based approach was assessed by classifying the sequences according to their contribution to the final taxonomic ID (Table 2).

**Table 2.**
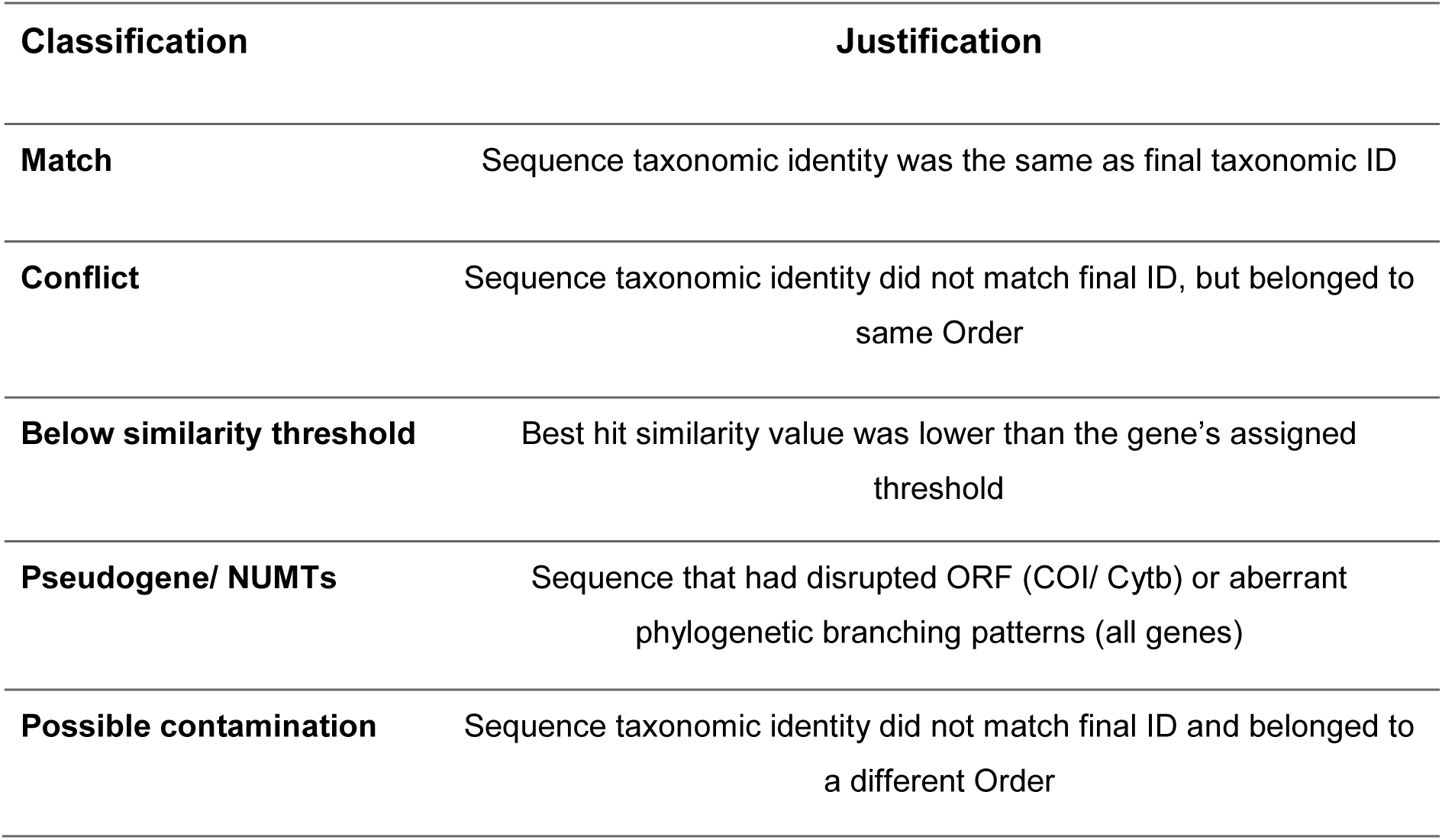
Sequence classification according to their contribution to the final identification of bushmeat samples.

| Classification | Justification |
| --- | --- |
| Match | Sequence taxonomic identity was the same as final taxonomic ID |
| Conflict | Sequence taxonomic identity did not match final ID, but belonged to same Order |
| Below similarity threshold | Best hit similarity value was lower than the gene's assigned threshold |
| Pseudogene/ NUMTs | Sequence that had disrupted ORF (COI/ Cytb) or aberrant phylogenetic branching patterns (all genes) |
| Possible contamination | Sequence taxonomic identity did not match final ID and belonged to a different Order |

An overall percentage of “lost” information was calculated among samples for all the available sequences (*N* = 8,692), considering sequences labelled as “Pseudogene / NUMTS”, “Below similarity threshold”, “Possible contamination” and “Conflict”, plus the amount of null PCR amplifications per gene. This process enabled us to determine the proportions of PCR error (contaminations and pseudogenes) and database gap (below thresholds and conflict).

For the purpose of improving taxonomic assignment accuracy, we reevaluated the species assignment thresholds fixed by Gaubert et al. (2015) using our incremented dataset. First, all the sequences matching with best hit similarity values <99% in DNAbushmeat, were re-blasted on NCBI. If blasting in NCBI resulted in a best match with the same species but with higher similarity percentage, the result was scored as an “improvement”. If the blasting in NCBI resulted in a best match with a different species, with higher similarity percentage, the result was scored as a “correction”. Second, we used the reassessed blast outcomes to establish new, empirical species assignment thresholds for each gene, considering that assignment error rate could not be over 10%. Because strict cut-offs do not fit with the complexity of DNA-based species assignment Final taxonomic IDs were updated on the basis of the corrections obtained from this re-blasting.

The contribution of DNA-typing in taxonomic assignment of bushmeat species was assessed in comparison with the original field ID conducted on the ground. We considered the final ID an “improvement” if it was included within the upper taxonomic rank resulting from the morphological ID (e.g., from *Cercopithecus sp.* or Primates to *Cercopithecus pogonias*). The molecular ID was considered a “correction” if it was not within the taxonomic range given by the morphological ID (e.g., *Cercopithecus campbelli* or Bovidae to *Cercopithecus mitis*).

### Macroecology of bushmeat market communities

The primary goal of using macroecological indices is to gain global insights into patterns and dynamics among bushmeat markets. Indeed, bushmeat marketplaces can be considered artificial communities of species that occur together in space and time, where interaction levels are conditioned by human activities (hunting, provisioning, selling, buying). Moreover, because bushmeat market communities are generally expected to constitute a representative proxy of the medium-to-large fauna present in the region (Gossé et al., 2022), assessing diversity patterns among markets may be a useful approach to characterise regional specificities in the trade.

Because of the acknowledged bias in sampling effort across countries, we did not use indicators relying on abundance (e.g., ɑ-diversity) and removed Gabon and Togo from the analyses (low sampling effort: *N* = 13-28). In order to identify hotspots of the bushmeat trade on a country basis and using samples’ final IDs, we estimated (i) species richness and (ii) phylogenetic diversity (PD) as a phylogenetic generalisation of species richness (Chao et al., 2010). By providing evolutionary context, PD is complementary to richness estimates as it can potentially uncover markets that impact species with crucial roles in ecosystem functioning due to their unique evolutionary traits. PD was calculated using the Faith Index (FI) (Faith, 1992). One representative sample per species was selected for final alignments. Domestic species (including *Bos taurus, Capra hircus, Ovis aries, Sus scrofa, Cavia porcella* and *Canis lupus familiaris*) were removed, as well as reptiles and birds due to uneven sampling among bushmeat markets. Phylogenetic trees were generated for each country from a four-gene concatenated matrix using BEAST2 v2.7.4 (Bouckaert et al., 2019). The Bayesian analysis was run under the bModelTest model with empirical frequencies, optimised relaxed clock and a Birth-Death model, for 50 million MCMC iterations.

Convergence of parameters was verified in Tracer 1.7.2

Phylogenetic β-diversity (PBD; (Graham & Fine, 2008) was estimated among countries in order to assess (i) the usefulness of this indicator to characterise the global dynamics of the bushmeat trade across tropical Africa and (ii) whether bushmeat market communities may be considered as representative proxies of regional biodiversity. We partitioned PBD into two components: phylogenetic turnover (changes in phylogenetic composition) and nestedness (level of shared phylogenetic history) (Baselga, 2009). Analyses were conducted in RStudio using the packages Picante for importing the phylogenetic trees and Betapart to calculate the PBD values. The phylogenetic tree was generated with the above-mentioned parameters (see Fig. S28, supplementary material).

Dissimilarity among bushmeat markets was visualised through non-metric multidimensional scaling (NMDS) using Jaccard dissimilarity matrices, with the Vegan package in RStudio. We used PERMANOVA (Anderson, 2008) through the Adonis function to test for significant differences in market composition. Homogeneity of dispersion (variance) within markets was estimated with the betadisper function (Anderson, 2006), and an ANOVA-like test (permutest) was used to determine if variances differed by markets.

We assessed the correlation between PBD and geographical distances between countries to further explore regional signatures in bushmeat trade biodiversity patterns. Geographic distances were calculated based on the shortest distance between country borders. Correlation was estimated through the Pearson correlation coefficient, including or excluding Benin as a potential outlier (see Results).

## Results

Overall, a total of 8,673 sequences were obtained from 2,487 samples, representing a sequencing success of 86.4%. Within the newly sequenced samples, COI had the highest rate of PCR failure (19.6%), on average 2.3 times greater than that of the other genes, with highest values for Pholidota, Carnivora and Cetartiodactyla (Table S2 and Fig. S29, Supplementary information). The mammalian orders that were most impacted by the amplification of potential pseudogenes were Pholidota (3.51% for 12S and 2.65% for Cytb) and Cetartiodactyla (3.34% for 12S). Regarding contamination levels, COI showed the highest values (12.90% for Pholidota). Proportions of best hit similarities below the species assignment threshold were similar across all genes, with overall values ranging between 3.22% and 3.33%. Primates were the most affected order (9.57%). Carnivora was the order with the highest level of conflict among sequence assignments, with 22.61% and 16.81% for 12S and 16S respectively (Fig. S30, Supplementary information).

Overall, the four-gene DNA-typing framework was capable of assigning species-level identifications to 96.07% (2516) of bushmeat samples, delineating a total of 133 validated species. A total of 84 samples could only be identified to the genus-level (20 distinct genera), five samples to the family-level (four Herpestidae and one Sciuridae) and nine samples to the order level (Cetartiodactyla, Primates and Rodentia). Among those, 34 taxa affiliated to 21 genera (including nine families: Bovidae, Canidae, Cercopithecidae, Herpestidae, Nesomyidae, Pythonidae, Sciuridae, Varanidae, Viperidae) could not be assigned to the species level, either because of cryptic diversity or gaps in reference databases (see Discussion and Table S1).

DNAbushmeat contributed 62.90% of the best-hit matches for COI, 63.50% for Cytb, 68.70% for 16S and 67.33% for 12S. Considering PCR failures, pseudogenes, contaminations, best hits inferior to similarity thresholds and gene conflicts, a single gene approach would have failed to reach final ID for 33.16% (COI), 24.02% (12S), 19.60% (Cytb) and 19.34% (16S) of the samples. Using a two-gene approach would have failed to reach final ID in c. 18% (Cytb-COI) to 26% (COI-12S) of the samples.

DNA-typing allowed for the refinement of 1,288 bushmeat carcasses, representing 54.30% of all the samples (Fig. 2). Corrections and improvements accounted for 569 (44.18%) and 719 (55.82%) of the refined morphological IDs. Within the most represented orders, Carnivora, Cetartiodactyla, Primates and Rodentia had the highest rates of correction (23.18-25.86%), while Pholidota and Primates showed the greatest levels of improvement (33.61-38.86%). Levels of refinement varied among countries, with the highest percentages obtained from European seizures at Roissy (France) and Zaventem (Brussels, Belgium) airports 95.08% and 92.08%, respectively.

**Figure 2.**
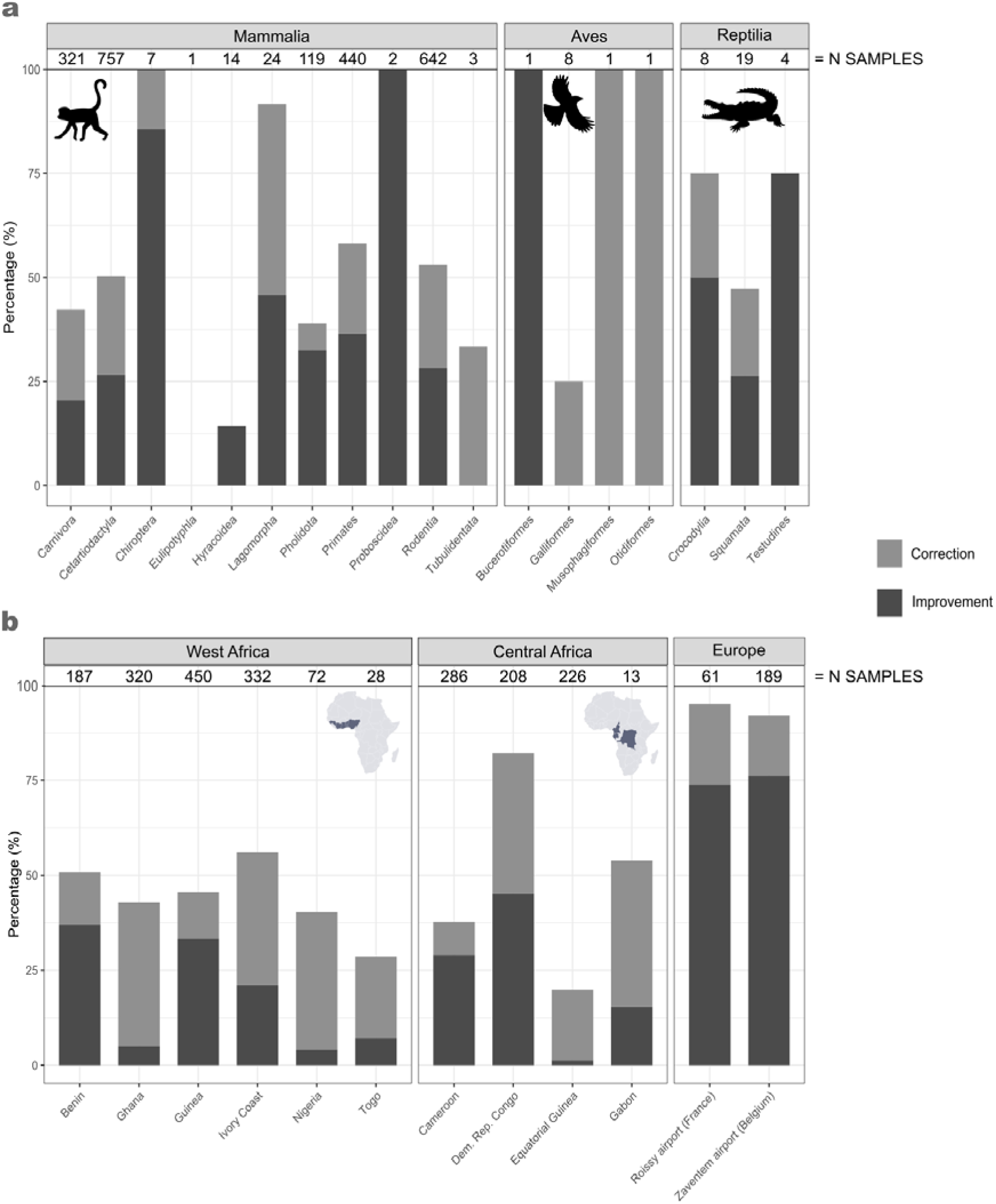
DNA-typing refinement of original field identifications across bushmeat taxa (a) and sampling origins (b).

Re-blasting all the sequences with best hit similarity values <99% obtained in DNA bushmeat on NCBI, 110 sequences had their best hit scores improved or were taxonomically reassigned for both 12S (10.60%) and 16S (10.62%). However, this was the case for only 37 (3.33%) and 50 (4.89%) sequences in COI and Cytb, respectively. On this basis and allowing for ≤10% assignment error rate (including best hit score improvement and taxonomic reassignment), species assignment thresholds were reassessed for the four genes, reaching 98.50% for COI, Cytb and 16S (grey zone = 98.00 - 99.00%), and 98.75% for 12S (grey zone 98.25 - 99.25%). For the fixed thresholds, empirically estimated error rates ranged from 8.11% (COI) to 9.09% (12S) and 10.00% (Cytb and 16S). At the terminal right distribution of the grey zone, error rates reached 0% (COI and Cytb), 0.91% (16S) and 1.92% (12S).

Final species ID detected 45 species and one genus (*Loxodonta* sp.) -representing 33.08% of the taxa identified- with a concerning conservation status (from CRitically Endangered to Near Threatened) sold in bushmeat markets from tropical Africa and trafficked into Europe. Primates, with a total of 23 species, were the most represented order, and we detected the greatest number of bushmeat species (18) of conservation concern from Equatorial Guinea (Fig. 3a and Fig. S31-S32, supplementary material). Species richness was the lowest in Nigeria with only 15 species, while Guinea was the highest, with a total of 52 species (Fig. 3b). The greatest values of phylogenetic diversity (PD) were found in the markets of Benin (6.1) and Guinea (5.8), whereas Ghana and European airports showed the lowest values (1.8-2.2) (Fig. 3c). Phylogenetic β-diversity (PBD) was the highest between Benin and DR Congo (0.81), with a strong phylogenetic turnover (0.71). PBD was the lowest between Ghana and Nigeria (0.39). The highest turnover was observed between DR Congo and Nigeria (0.73), while the lowest turnover was between Ghana and Ivory Coast (0.02). Phylogenetic nestedness was the highest between Guinea and Nigeria (0.51) and the lowest between Cameroon and Equatorial Guinea (0.013) (Table 3).

**Figure 3.**
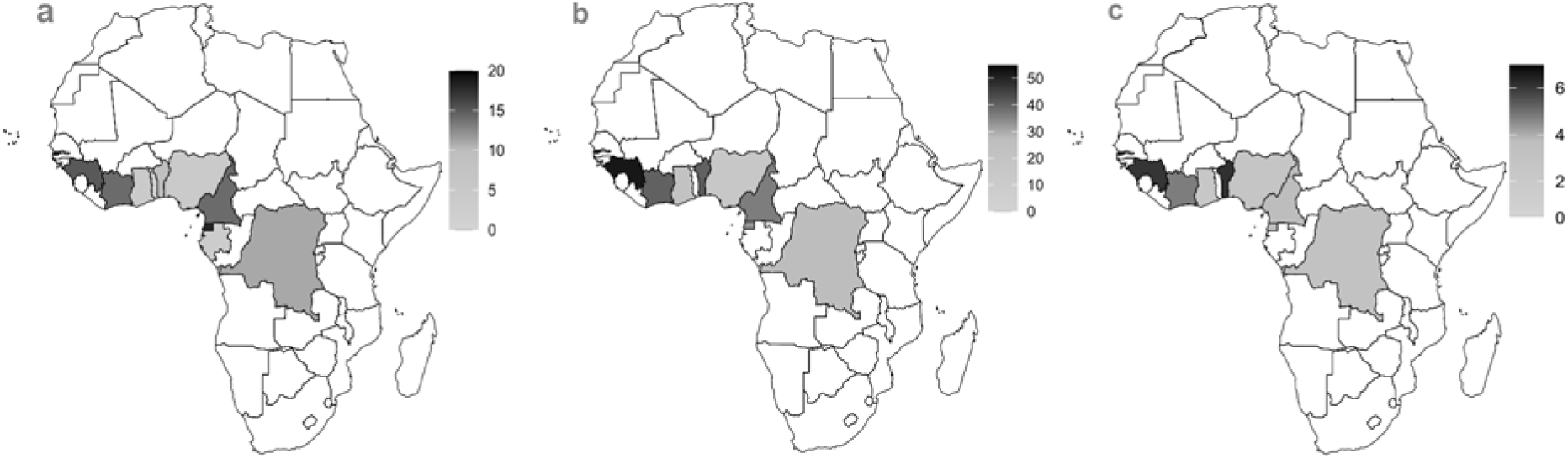
Conservation status and richness profiles of the African bushmeat taxa at the scale of national markets. a) Number of species of conservation concern according to the IUCN Red List of Threatened Species (from Near Threatened to Critically Endangered); b) Species richness (N species / taxa); c) Phylogenetic diversity (Faith Index). Gabon and Togo were excluded from maps (b) and (c) because of insufficient sampling efforts. Maps were created with package *rnaturalearth* in RStudio.

**Table 3.**
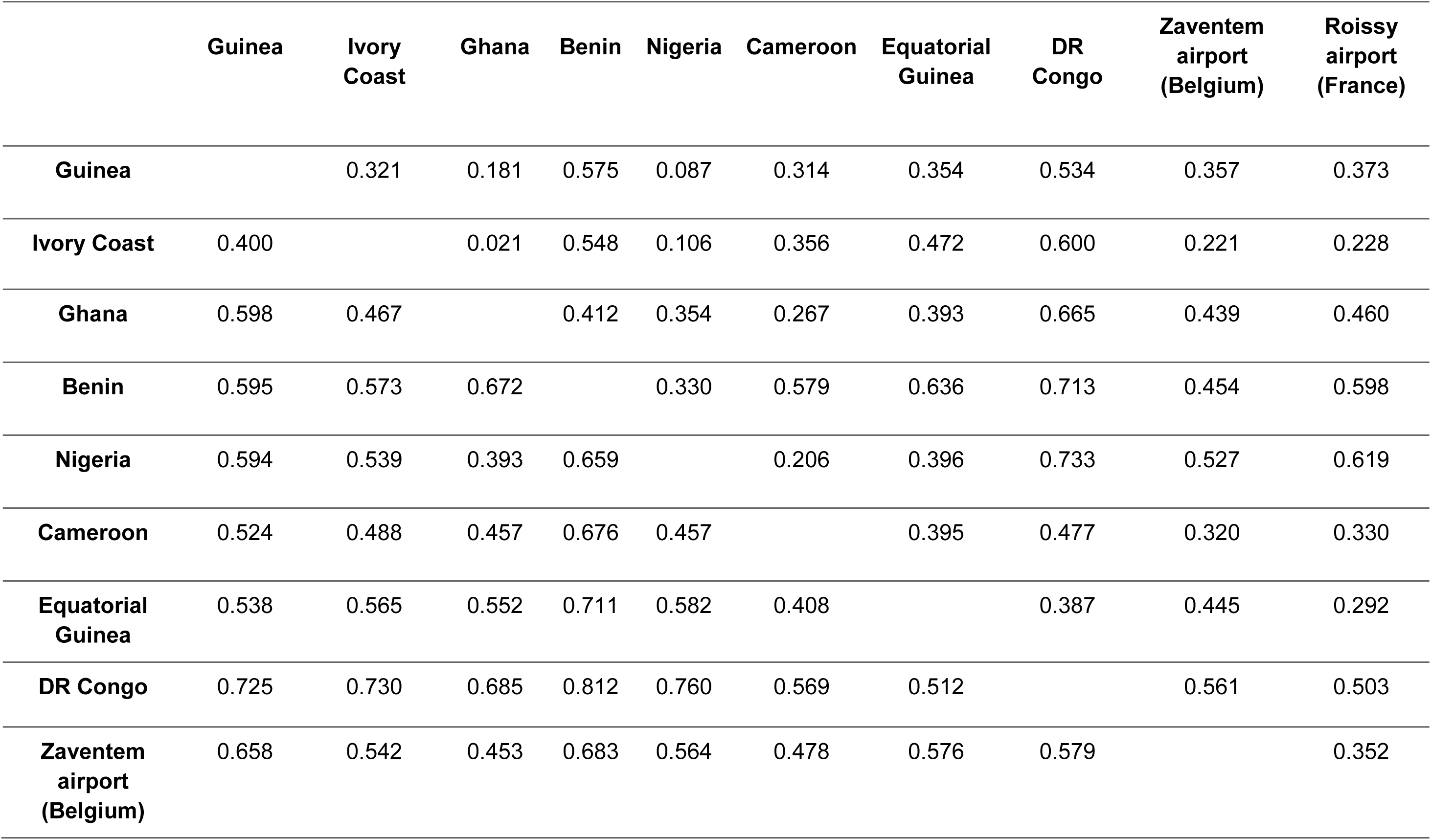
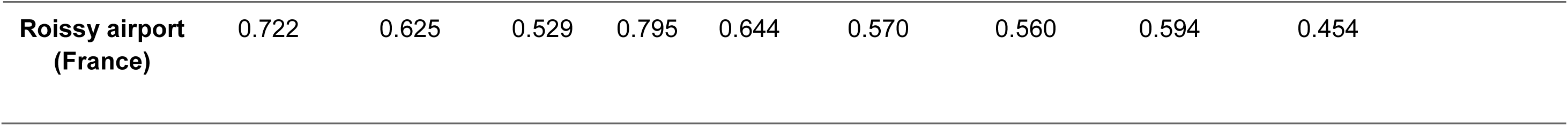
Distribution of phylogenetic β-diversity (below empty diagonal) and turnover (above empty diagonal) between countries.

NMDS analysis revealed five distinct regional clusters among national bushmeat markets that correspond to the following biogeographic regions: Upper Guinean Block (Guinea and Ivory Coast), Dahomey Gap (Benin), Lower Guinean Block (Cameroon and Equatorial Guinea), and Congolia (DR Congo). Ghana (covering the Upper Guinean Block and Dahomey Gap) and SW Nigeria (Dahomey Gap) constituted the fifth cluster. PERMANOVA indicated statistically significant differences between markets (*p*=0.045) with non-significant differences in dispersion within markets (ANOVA; *p*=0.124).

Correlation between PBD and geographic distances between countries was moderate (*r*=, 0.47) when Benin was included. In this case, pairwise comparisons with Benin were always above the trend line, neighbouring countries (e.g. Nigeria) included, in line with an outlier pattern. After removing Benin, correlation between PBD and geographic distances was strong (*r*=0.63) (Fig. 5; also see Fig. S33, supplementary material). Nigeria, as the remaining representative of Dahomey Gap, generally maintained pairwise comparisons above the trend line.

**Figure 4.**
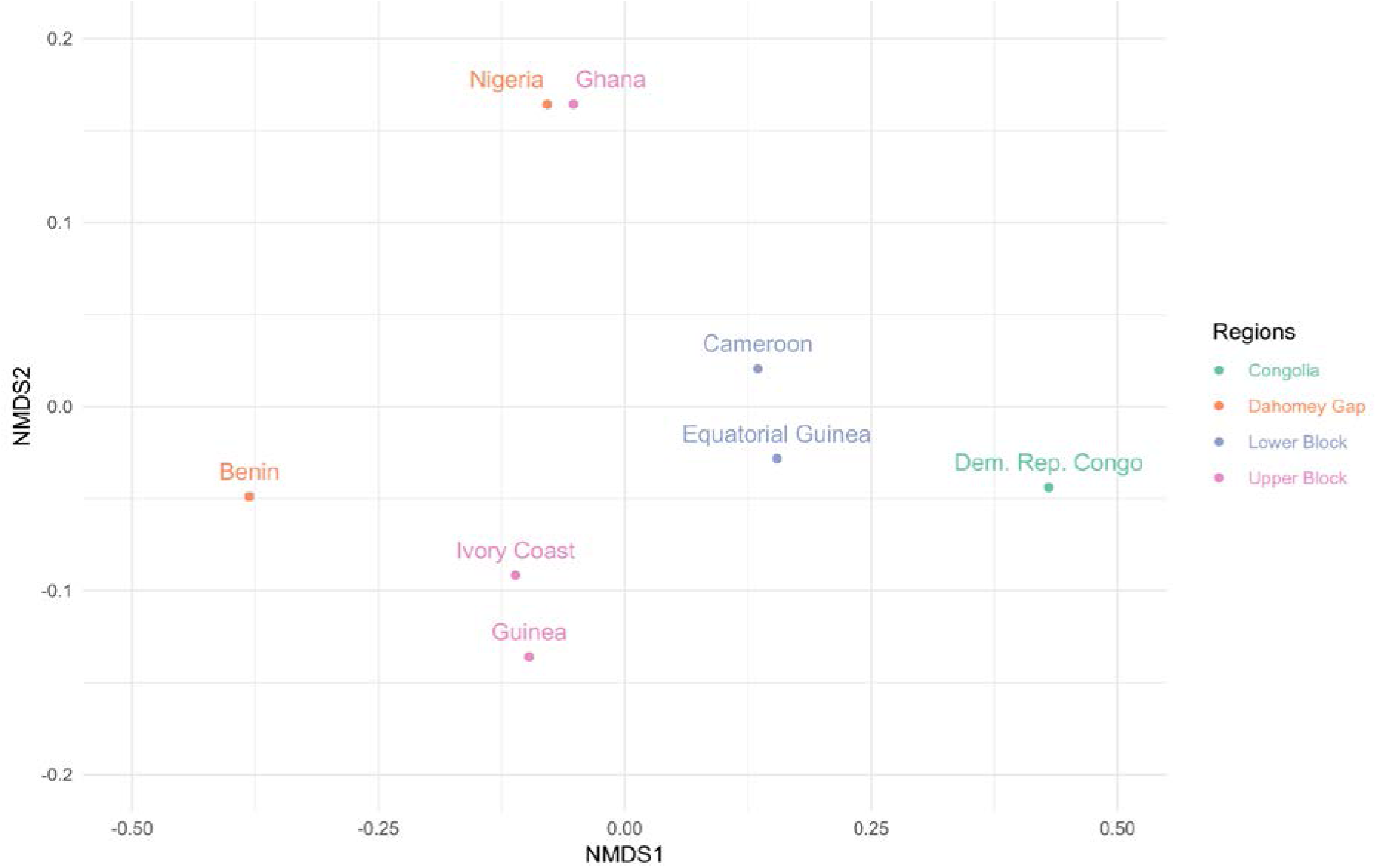
Non-Metric Multidimensional Scaling (NMDS) plot of national bushmeat markets according to phylogenetic β-diversity. Bushmeat markets are grouped by biogeographic ecoregions across tropical Africa. Airport seizures, as well as Gabon and Togo were excluded due to admixed sources and low sampling efforts, respectively.

**Figure 5.**
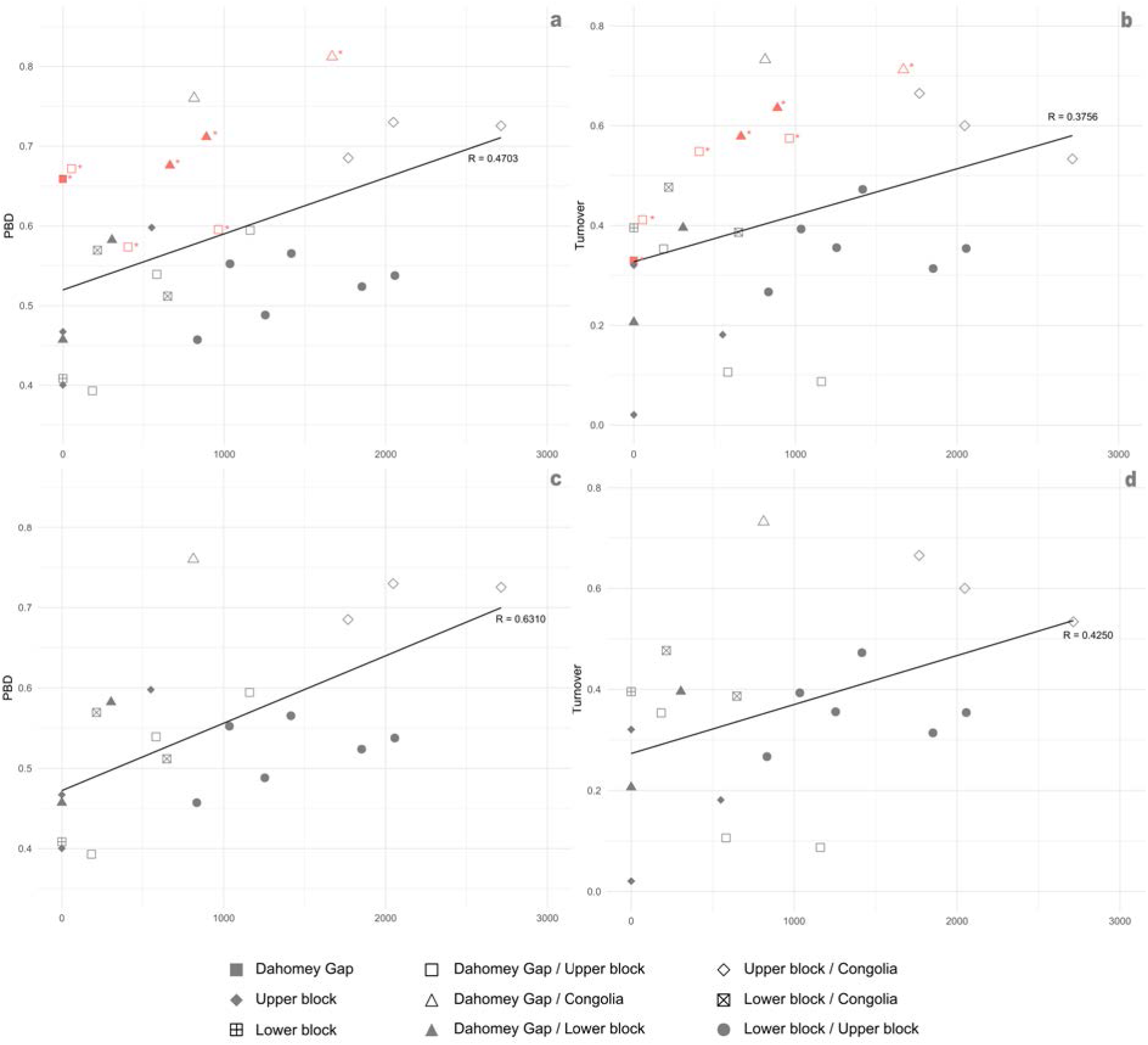
Correlation trends between national bushmeat markets’ phylogenetic β-diversity (left) and phylogenetic turnover (right), and distances between countries. (a-b) including Benin; (c-d) excluding Benin.

## Discussion

### Four-gene DNA-typing approach significantly improves bushmeat species identification

Our study represents an unprecedented genetic survey of the bushmeat trade in tropical Africa and Europe, totalling 8,673 mitochondrial sequences –of which more than half has been newly generated– from 2,487 samples representing 133 validated species. Since the first efforts to produce a four-gene reference database aimed at tracing the bushmeat trade (Gaubert et al., 2015), the number of sequences generated, the number of DNA-typed bushmeat samples and the number of species surveyed were multiplied more than seven, eight and two times, respectively. Such wide coverage answers the call for large scale studies capable of better apprehending the dynamics of the bushmeat trade, and contrasts with the relatively lower sequencing efforts (50-350 samples) from previous bushmeat genetic studies (Groom et al., 2023).

Our study further demonstrates that bushmeat DNA-typing is affected by biases variously distributed across genes and taxa, including PCR failure, pseudogene amplification, sequencing of exogenous (non-target) DNA, and incomplete DNA registers. Although mtDNA has long been the most prominent marker for assessing species identity within vertebrates (Elyasigorji et al., 2022), the genetic description of African vertebrate diversity -which seems affected by unsuspected levels of cryptic diversity-remains incomplete (e.g., Gossé et al., 2022; Mulvaney et al., 2023). Moreover, the non-optimal quality of bushmeat samples, which have often gone through post-mortem tissue degradation and meat processing, and potential cross-species DNA contamination occurring on the market stalls, are factors likely promoting the above-mentioned PCR biases (Gaubert et al., 2015). COI, though widely accepted as a universal marker for animal species identification (Ahmed et al., 2022), provided the worst outcomes of the four genes, with a PCR failure rate 2.3 times higher than that of other mtDNA genes. Nearly 30% of the samples representing taxa frequently sold as bushmeat (Pholidota, Carnivora, Cetartiodactyla, Primates, and Rodentia) were affected by such PCR failure. This can be explained by the non-optimal design of COI universal primers due to the continuous distribution of polymorphism along the gene, resulting in the highly degenerate nature of COI primers (see Gaubert et al., 2015).

Based on the empirical evidence provided as part of this study, we advocate for the use of a four-gene approach in order to reach accurate bushmeat species identification. Indeed, biases affecting DNA-typing are likely to be tempered by a multigene approach (including COI), involving different primer properties and PCR sensitivity, and querying across different DNA registers. In our case, a single-gene approach would have led to inconclusive or erroneous species identification in c. ¼ (Cytb, 12S, 16S) to ⅓ (COI) of the sample set. We further show that a two-gene approach, despite being commonly used in bushmeat surveys (Deagle et al., 2014; Gombeer et al., 2020), would not be sufficient to reach species identification in c. 18-26% of the cases.

Overall, our four-gene DNA-typing framework was able to assign final species-level identification to c. 96% of the bushmeat samples, which is in line with the performance of recent surveys using the same protocol (Din Dipita et al., 2022; Gaubert et al., 2015; Gossé et al., 2022). Most of the remaining samples (3.3%) were identified to the genus-level. One of the main reasons involved is incomplete DNA registers, where species have not yet been DNA-typed (e.g., within *Kobus*, Sciuridae and Cercopithecidae), together with incomplete knowledge on species range (e.g., *Colobus*). Another important factor relates to the limitation of using a single locus (mtDNA) in the framework of complex evolutionary processes involving cryptic diversity (e.g., *Crossarchus*), super-species and introgression (e.g., *Genetta*, *Loxodonta*, Cercopithecidae). Although the total proportion of samples not reaching species level identification remains low (c. 4%), further recommendations to improve the DNA-typing procedure could include sequencing approaches allowing full mitogenome assembly (improving resolution among all taxa but limited to a single locus) and the use of rapidly evolving nuclear markers (improved resolution among targeted taxa).

One of the major challenges in traditional bushmeat surveys is to reach the correct morphological identification of carcasses (Schilling et al., 2020). The majority of animals are sold smoked and / or processed and butchered into various forms obscuring specific diagnostic traits (Minhós et al., 2013; Olayemi et al., 2011). Additional complexity may also arise from the use of local terminologies by sellers together with reliance on third parties as collectors, and in our case, the heterogeneity in field identification protocols, all yielding poor species identification hypotheses (Din Dipita et al., 2022; Gaubert et al., 2015). Overall, our results highlight the significant limitation of morphological identification and underscore the essential role of the four-gene DNA-typing approach in producing accurate identifications of bushmeat species. Indeed, more than half of the samples underwent taxonomic refinement through DNA-typing, of which c. 46% were taxonomic corrections (wrong morphological hypothesis). The efficiency of our molecular assignment protocol may explain the higher rate of refinement that we obtained compared to other studies (c. 19-30%; Among some of the most represented mammalian orders in the market (Carnivora, Cetartiodactyla, Pholidota, Primates and Rodentia), the accuracy of morphological identification varied from c. 38 to 60 %. Primates showed the highest rates of misidentification (c. 62%), in line with previous studies from Cameroon and Equatorial Guinea, and due to their complex evolutionary history and the post-processing removal of their species-diagnostic phenotypic traits (Din Dipita et al., 2022; Minhós et al., 2013). Levels of taxonomic refinement were variable among African countries and reached 82.2 % in the case of DR Congo. However, the most striking contributions of DNA-typing were observed from European seizures at the airports of Roissy (France; 95% of refinements) and Zaventem (Belgium; 92%). European airports have recently been pinpointed as important hubs of the bushmeat trade from Africa (Chaber et al., 2010; Falk et al., 2013), and accurate identification of the bushmeat species is a crucial prerequisite to illegal trade mitigation and law enforcement (Chaber et al., 2023; Morrison-Lanjouw et al., 2023). In the case of airports, refinement was mostly driven by increased taxonomic resolution (improvements rather than corrections), exemplifying the challenge faced by customs to reach accurate species identification from the highly processed carcasses transported in customers’ luggage (Chaber et al., 2023; Din Dipita et al., 2022).

The various constraints inherent to bushmeat surveys (expert knowledge requisite, vertebrates’ cryptic diversity, meat processing) further complicate the morphological identification of carcasses and are a general limitation to the reliability of traditional survey methods (Din Dipita et al., 2022; Djagoun et al., 2023; Gossé et al., 2022). The four-gene DNA typing approach that we apply addresses these limitations by offering consistent, objective and reliable identification of bushmeat carcasses, regardless of their condition and the identification protocols applied in the field.

### Optimising bushmeat taxonomic assignment with empirically reassessed species thresholds and expert referencing in Genbank

The delineation of genetic species thresholds is a fundamental challenge in molecular taxonomy (Hebert et al., 2003; Twyford et al., 2024; Waugh, 2007). Establishing such thresholds is critical, as they directly influence taxonomic resolution accuracy and the ability to detect cryptic biodiversity—one of the key advantages of DNA-based approaches over traditional morphological methods (Packer et al., 2009). However, defining these thresholds is inherently complex, as they are case-dependent, varying with the genes and species being studied. Consequently, threshold values derived from empirical evidence have been recommended

At the time of constructing DNAbushmeat, a fixed, conservative threshold of ≥95% similarity was applied to minimize potential assignment bias resulting from the partial taxonomic representation (60 species) of the database. Here, we follow original recommendations advocating for an empirical reassessment of this threshold (Gaubert et al., 2015), leveraging the expanded taxonomic representation now available (133 species). By re-blasting all sequences with initial best-hit similarity values below 99% in DNAbushmeat, we redefined species-level thresholds specific to African bushmeat species and the levels of polymorphism of the four genes analyzed. The revised thresholds were determined as 98.50% for COI, Cytb, and 16S, and 98.75% for 12S. These empirically derived thresholds are now accompanied by a 1% ‘grey zone’ to mitigate the risks of false positives and false negatives associated with fixed thresholds (Zhang et al., 2017). This provides a biologically meaningful framework for species hypothesis testing, better fitting the potential evolutionary complexity of the studied species. Specifically, best-hit similarity values within the +0.5% range above the species threshold should be considered highly probable species-level assignments, whereas values within the -0.5% range below the threshold should require further validation using additional lines of evidence.

Accurate species identification is a crucial prerequisite for effective conservation efforts (Francis et al., 2010). The refined genetic species thresholds we propose will enhance the efficiency of bushmeat trade surveys, with direct implications for conservation. DNA-typing enabled the identification of approximately one-third of the vertebrate taxa sold in tropical African bushmeat markets as species of conservation concern, according to the IUCN Red List of Threatened Species. However, inaccurate taxonomic identifications—often occurring as carcasses are butchered or processed—can hinder efforts to monitor and protect species at risk of extinction (Deagle et al., 2014). Furthermore, since taxa identified as conservation priorities are frequently the target of illegal trafficking (e.g., pangolins, Heinrich et al., 2016), precise species identification is essential for strengthening law enforcement and ensuring compliance with wildlife protection regulations. Primates, with a total of 23 species, were the most represented order, exhibiting both a predominant presence in market stalls across central Africa (namely, Cameroon and Equatorial Guinea) and high conservation concern statuses (Fernandez et al., 2022). In regions where bushmeat trade significantly affects vulnerable species—such as Equatorial Guinea, where 18 species (c. 49% of the total recorded in the country) were identified—reliable identification tools can improve monitoring efforts and optimize resource allocation diversity. Overlooking taxonomic diversity can have important consequences for ecosystem stability, conservation strategies, and even disease control (Hending, 2025). Traded African mammals and other terrestrial vertebrates likely harbour substantial but previously unrecognized cryptic diversity (e.g., Din Dipita et al., 2022; Eaton et al., 2009; Gaubert et al., 2015; Gossé et al., 2022), sometimes leading to new species descriptions (Colyn et al., 2010; Oates et al., 2022). In our study, extensive bushmeat screening combined with rigorous reference database verification for the relevant genes led to the identification of two potentially undescribed mongoose species (Herpestidae) from West Africa (Guinea, Togo and Benin). Indeed, 17 samples associated with these taxa yielded best-hit values below the newly established thresholds –including the delimited grey zone range (92.1(COI) - 98.7(16S)%), and their phylogenetic uniqueness was confirmed through phylogenetic tree analysis (data not shown). Applying the same methodology, we also detected a certain level of cryptic diversity, defined here as bushmeat taxa with best-hit values falling within the lower distribution of the grey zone (98.28 (COI) - 100 (12S)%). Notably, this included 14 samples of guenon (*Cercopithecus* spp.) from Cameroon, affiliated with the *Cercopithecus mona* supergroup (Kingdon et al., 2013). Lastly, some cases remained unresolved, where taxonomic gaps in reference databases could not be distinguished from cryptic or undescribed diversity. This was particularly evident in 11 samples of squirrels from the Protoxerini tribe, representing seven distinct but unassigned species. Of the 30 described Protoxerini species (Thorington Jr. & Hoffmann, 2005), only seven were represented in Genbank.

Since DNAbushmeat contributed to only two-thirds of the best-hit matches (relative to NCBI) and is no longer updated or operational, here we provide guidelines for applying the species-assignment methodological assay and the bushmeat database developed in this study directly on NCBI. All sequences published and analyzed (N= 8692 sequences) in this work have been tagged with the descriptor |collected_by="DNAbushmeat"| in Genbank. To achieve taxonomic identification of a given bushmeat sequence using our expert reference database, users should perform a BLAST search on the blastn webpage of the NCBI platform (NCBI BLAST). After inputting the sequence(s) or a sequence file (e.g., FASTA format), users should refine the search to the DNAbushmeat nucleotide database by specifying |collected_by="DNAbushmeat"| in the Entrez Query field within the Choose Search Set section. We recommend using Megablast (Zhang et al., 2000) to optimize the search for closely related sequences, with default algorithm parameters (including 100 maximum best hits and 0.05 as expect threshold). The BLAST results can then be downloaded and further analyzed using the species-level assignment thresholds outlined in this study.

### Leveraging community ecology indexes for large-scale surveillance of the bushmeat trade

Diversity indices have been widely used to identify large-scale biodiversity patterns, such as global hotspots (e.g., Myers, 1990; Sechrest et al., 2002), and to assess the impact of human activities on biodiversity (e.g., Socolar et al., 2016). More recently, these indices have been applied to the bushmeat trade to identify key trade hotspots that significantly affect the phylogenetic legacy of African vertebrate diversity (Gossé et al., 2022). However, several factors constrain bushmeat trade surveys, potentially biasing species richness and diversity indicators. Disparate sampling efforts across countries, driven by logistical constraints, can result in uneven taxonomic sampling (Sánchez-Fernández et al., 2022). This was the case for Gabon and Togo (*N* = 13-28), which are therefore not considered in this section. Another source of bias lies in the nature of the bushmeat sites surveyed. Large urban markets typically source from a broader geographic range and a wider variety of taxa (McNamara et al., 2016), whereas smaller rural markets tend to rely on nearby forests, leading to reduced taxonomic diversity (Sackey et al., 2022). Similarly, mid-sized markets in regions with naturally low faunal diversity tend to offer a more limited selection of species. This pattern was observed in Nigeria, where only 15 species could be identified. Despite the country’s overall high biodiversity, the surveyed bushmeat market was located in the southwest (Asejire; see Olayemi et al., 2011), a region where lower mammalian diversity is expected (Booth, 1958).

On the other hand, Guinea recorded the highest species richness (52 species) and the second-highest phylogenetic diversity value (5.8). This reflects the presence of markets sourcing bushmeat from both forested regions in southeastern Guinea and forest-savannah habitats in the east, thereby maximizing species richness estimates. Despite being part of the Dahomey Gap—a region of West Africa considered to have relatively low biodiversity (Booth, 1958)—Benin unexpectedly exhibited the highest phylogenetic diversity (6.1), nearly three times that of southwestern Nigeria, which is also within the Dahomey Gap (Booth, 1958). This remarkable diversity, spanning 14 different taxonomic orders of terrestrial vertebrates, highlights Benin’s extensive wildlife trade network, where bushmeat and traditional medicine markets are closely interconnected. The latter, in particular, often sources species from a wider geographic range, including foreign countries (Djagoun et al., 2023).

Discrepancies in experimental design, which are inherent to any bushmeat survey (see Groom et al., 2023), were mitigated by maintaining a sampling strategy that ensured (i) unbiased taxonomic sampling (i.e., no preference for specific taxa) and (ii) coverage across diverse market types and geographic regions within each country. A notable exception was in DR Congo, where only two urban bushmeat markets—Kinshasa and Kisangani—and a narrower taxonomic spectrum were surveyed. Taking into account these dataset limitations and the potential impact of taxonomic sampling bias on phylogenetic diversity estimates (Park et al., 2018), we assessed the utility of phylogenetic β-diversity (PBD) to characterize the structural dynamics of national bushmeat markets. Our primary objective was to apply a macroecological approach to examine global diversity patterns in the bushmeat trade across tropical Africa and test whether bushmeat markets reliably reflect vertebrate diversity. Specifically, we assess whether national bushmeat market trends align with the expected biogeographic patterns shaping vertebrate communities from West to Central Africa (Droissart et al., 2018; White, 1979). This approach aims to provide a global view on the bushmeat trade in tropical Africa and inform future regional-scale strategies for trade regulation.

Notably, lower and higher phylogenetic β-diversity (PBD) values suggest high and low faunal assemblage similarity between national bushmeat markets, respectively. Higher PBD turnover indicates phylogenetically distinct communities, whereas higher PBD nestedness reflects disparities in species richness between markets. However, since nestedness is highly sensitive to sampling effort (Soininen et al., 2017), we will exclude it from our discussion. Our results indicate that, despite variations in survey design, national bushmeat markets can reliably reflect vertebrate diversity within states, particularly for rainforest species. Overall, global PBD and PBD turnover values were strongly correlated with geographic distances—an expected pattern given the west-to-east gradient (∼4000 km) in tropical Africa, which encompasses distinct biogeographic regions: the Upper Guinean Block (western), the Lower Guinean Block (central), and Congolia (eastern) (Hardy et al., 2013). NMDS analysis further supports this pattern, recovering three regional clusters corresponding to these biogeographic zones: Upper Guinean Block (Guinea and Ivory Coast), Lower Guinean Block (Cameroon and Equatorial Guinea), and Congolia (DR Congo). This differentiation highlights the critical role of ecoregions in shaping biodiversity distribution and trade dynamics, as ecological factors likely influence species composition and market trends within these regions (Petrozzi et al., 2016).

NMDS analysis also identified two additional clusters, differentiating Benin from Ghana and southwestern Nigeria, both groups encompassing the Dahomey Gap area as defined by zoologists (Moreau, 2009). Benin stood as a highly differentiated entity from the other African markets and an outlier in the PBD-geographic distance correlation trend, and was clearly demarcated from the rest of the bushmeat markets representative of the Dahomey Gap area. As Benin is a global hub for wildlife trade relying on a wide spectrum of sources (see above), it represents a major hotspot of the bushmeat trade, which our community ecology-based evaluation was able to successfully characterize. While we initially expected Ghana to cluster more closely with other Upper Guinean Block countries, it instead grouped with southwestern Nigeria, a biogeographic representative of the Dahomey Gap. Although much of Ghana’s forested areas are biogeographically assigned to the Upper Guinean Block, our findings suggest an alternative pattern influencing bushmeat sourcing for the country’s major markets—one that may favor the trade of ubiquitous species found in southeastern Ghana.

## Conclusion

Beyond the unprecedented coverage of bushmeat markets and the empirically refined DNA-based identification of bushmeat species in our study, further improvements in survey design can be foreseen. We posit that bushmeat markets function as ecological communities with relatively simple and constrained dynamics, where species diversity fluctuates over time through a birth-death process (i.e., animals entering and leaving the market via trade fluxes) but remains stable in space (see Gossé et al., 2022). Building on this premise, we argue that further integrating community ecology frameworks holds significant potential for uncovering the broader dynamics of the bushmeat trade. This, in turn, could facilitate the development of scale-adapted indicators to inform more effective management strategies for the bushmeat trade. Further sampling efforts are needed to expand the taxonomic scope of DNA-typing, improve the detection of cryptic diversity, and enhance the standardization of experimental design across study sites. Additionally, controlling for false negative rates (i.e., undetected species) will help refine comparative analyses and ensure more robust assessments. Our findings demonstrate that the proportion of IUCN Red Listed species, along with measures of species richness, phylogenetic diversity, and phylogenetic β-diversity, serve as valuable indicators for characterizing the structural patterns of the bushmeat trade. A geographically informed approach within defined ecoregions of tropical Africa could strengthen transboundary coordination in conservation efforts (Groom et al., 2023). These indicators were particularly instrumental in highlighting the scale of wildlife trade in Benin, emphasizing the urgent need for mitigation at two key levels: preserving local ecosystems and improving cross-border trade management.

## Acknowledgments

The genetic analyses were conducted from 2010 to 2023 in three different laboratories, including Service de Systématique Moléculaire of the Muséum National d’Histoire Naturelle in Paris (MNHN), Institut des Sciences de l’Evolution de Montpellier (ISEM) at Université de Montpellier, and Centre de Recherche sur la Biodiversité et l’Environnement (CRBE) at Université de Toulouse. We thank the staff from those molecular platforms for continuous support and fruitful discussions on experimental design, including C. Bonillo and J. Lambourdière at MNHN, Fabienne Justy and Frédérique Cerqueira at ISEM, and the B2M team at CRBE. Joseph Brennan, Jena McKelvey, Monica Youk, Nicolas Leroy, Antoine Signol, and Damien De Craieye have contributed to the sequencing effort. We thank Remy Dernat (Institut des Sciences de l’Evolution de Montpellier) for technical assistance with a remanence of DNABUSHMEAT. We gratefully acknowledge the following individuals for their contributions to the sampling effort: Paolo Pagani (Utrecht University), Jules Koffi Gossé (Université F. Houphouët-Boigny), Nutsuakor Mac Elikem (Kwame Nkrumah University of Science and Technology, Kumasi), Stanislas Zanvo (Université d’Abomey Calavi), Flobert Njiokou (Université de Yaoundé I), Alain Din Dipita (Université de Douala), Pablo Esono Esono (ANDEGE), Idriss Ayaya (Université de Kisangani), Jonas Muhindo (Solutions for Wildlife / Université de Kisangani), V. Benoît and L. Cervantes (Institut de Recherche Criminelle de la Gendarmerie Nationale, Cergy Pontoise, France), and Véronique Renault (Université de Liège, Belgium).

Sampling effort and lab work were funded by Action Transversale Muséum (MNHN), Société des Amis du Muséum National d’Histoire Naturelle et du Jardin des Plantes, Consortium National de Recherche en Génomique, @SPEED-ID (Accurate SPEcies Delimitation and Identification of eukaryotic biodiversity using DNA markers’ of the French Barcode of life Initiative), Fundação para a Ciência e a Tecnologia (FCT IC&DT 02/SAICT/2017—n◦ 032130; BUSHRISK), Strategic Funding U-IDB/04423/2020 and UIDP/04423/2020 through national funds provided by the FCT and the European Regional Development Fund (ERDF) in the framework of the program PT2020, European Structural and Investment Funds (ESIF) through the Competitiveness and Internationalization Operational Program - COMPETE 2020, Agence Nationale de la Recherche (ANR-17-CE02-0001; PANGO-GO), Institut de Recherche pour le Développement (Jeune Equipe Associée à l’IRD; RADAR-BE), Ministère de l’Enseignement Supérieur et de la Recherche Scientifique of Côte d’Ivoire (C2Ds-IRD; TRACE-BROUSSE), and the Belgian Federal Public Service Health, Food Chain Safety and Environment (Cahier spécial des charges n° DG5/AMSZ/LF/16018).

## Author Contributions

**Daniel Pires:** Data curation, formal analysis, Writing - original draft, Writing - review & editing. **Cidalia Gomes:** Data curation, Writing - review & editing. **Belinda Groom:** Data curation, Writing - review & editing. **Sylvain Dufour:** Resources, Writing - review & editing. **Emmanuel Danquah:** Resources, Writing - review & editing. **Nathalie Van Vliet:** Resources, Writing - review & editing. **Gabrial Ngua Ayecaba:** Resources, Writing - review & editing. **Alain Didier Missoup:** Resources, Writing - review & editing. **Komlan Afiademanyo:** Resources, Writing - review & editing. **Chabi Djagoun:** Resources, Writing - review & editing. **Sery Bi Gonedele:** Resources, Writing - review & editing. **Anne-Lise Chaber:** Resources, Writing - review & editing. **Ayodeji Olayemi:** Resources, Writing - review & editing. **Agostinho Antunes:** Conceptualization, Funding acquisition, Investigation, Methodology, Project administration, Resources, Supervision, Validation, Writing – review & editing. **Philippe Gaubert:** Conceptualization, Data curation, Formal analysis, Funding acquisition, Investigation, Methodology, Project administration, Resources, Supervision, Validation, Writing – original draft, Writing – review & editing.

