## Supplementary Material for "Advancing global DNA-referencing of bushmeat in African tropical forests: a tool for identifying trade hotspots and dynamics in western and central Africa"

**Figure S1.** Optimal COI Neighbour Joining tree of Carnivora bushmeat samples.

**Figure S2.** Optimal COI Neighbour Joining tree of Carnivora bushmeat samples.

**Figure S3.** Optimal 16S Neighbour Joining tree of Carnivora bushmeat samples.

**Figure S4.** Optimal 12S Neighbour Joining tree of Carnivora bushmeat samples.

**Figure S5.** Optimal COI Neighbour Joining tree of Cetartiodactyla bushmeat samples.

**Figure S6.** Optimal Cytb Neighbour Joining tree of Cetartiodactyla bushmeat samples.

**Figure S7.** Optimal 16S Neighbour Joining tree of Cetartiodactyla bushmeat samples.

**Figure S8.** Optimal 12S Neighbour Joining tree of Cetartiodactyla bushmeat samples.

**Figure S9.** Optimal COI Neighbour Joining tree of Hyracoidea bushmeat samples.

**Figure S10.** Optimal 16S Neighbour Joining tree of Hyracoidea bushmeat samples.

**Figure S11.** Optimal 12S Neighbour Joining tree of Hyracoidea bushmeat samples.

**Figure S12.** Optimal COI Neighbour Joining tree of Lagomorpha bushmeat samples.

**Figure S13.** Optimal Cytb Neighbour Joining tree of Lagomorpha bushmeat samples.

**Figure S14.** Optimal 16S Neighbour Joining tree of Lagomorpha bushmeat samples.

**Figure S15.** Optimal 12S Neighbour Joining tree of Lagomorpha bushmeat samples.

**Figure S16.** Optimal COI Neighbour Joining tree of Pholidota bushmeat samples.

**Figure S17.** Optimal Cytb Neighbour Joining tree of Pholidota bushmeat samples.

**Figure S18.** Optimal 16S Neighbour Joining tree of Pholidota bushmeat samples.

**Figure S19.** Optimal 12S Neighbour Joining tree of Pholidota bushmeat samples.

**Figure S20.** Optimal COI Neighbour Joining tree of Primates bushmeat samples.

**Figure S21.** Optimal Cytb Neighbour Joining tree of Primates bushmeat samples.

**Figure S22.** Optimal 16S Neighbour Joining tree of Primates bushmeat samples.

**Figure S23.** Optimal 12S Neighbour Joining tree of Primates bushmeat samples.

**Figure S24.** Optimal COI Neighbour Joining tree of Primates bushmeat samples.

**Figure S25.** Optimal Cytb Neighbour Joining tree of Primates bushmeat samples.

**Figure S26.** Optimal 16S Neighbour Joining tree of Primates bushmeat samples.

**Figure S27.** Optimal 12S Neighbour Joining tree of Primates bushmeat samples.

**Figure S28.** Phylogenetic tree generated from representative samples of each species from a four-gene concatenated matrix with 100 000 000 MCMC iterations.

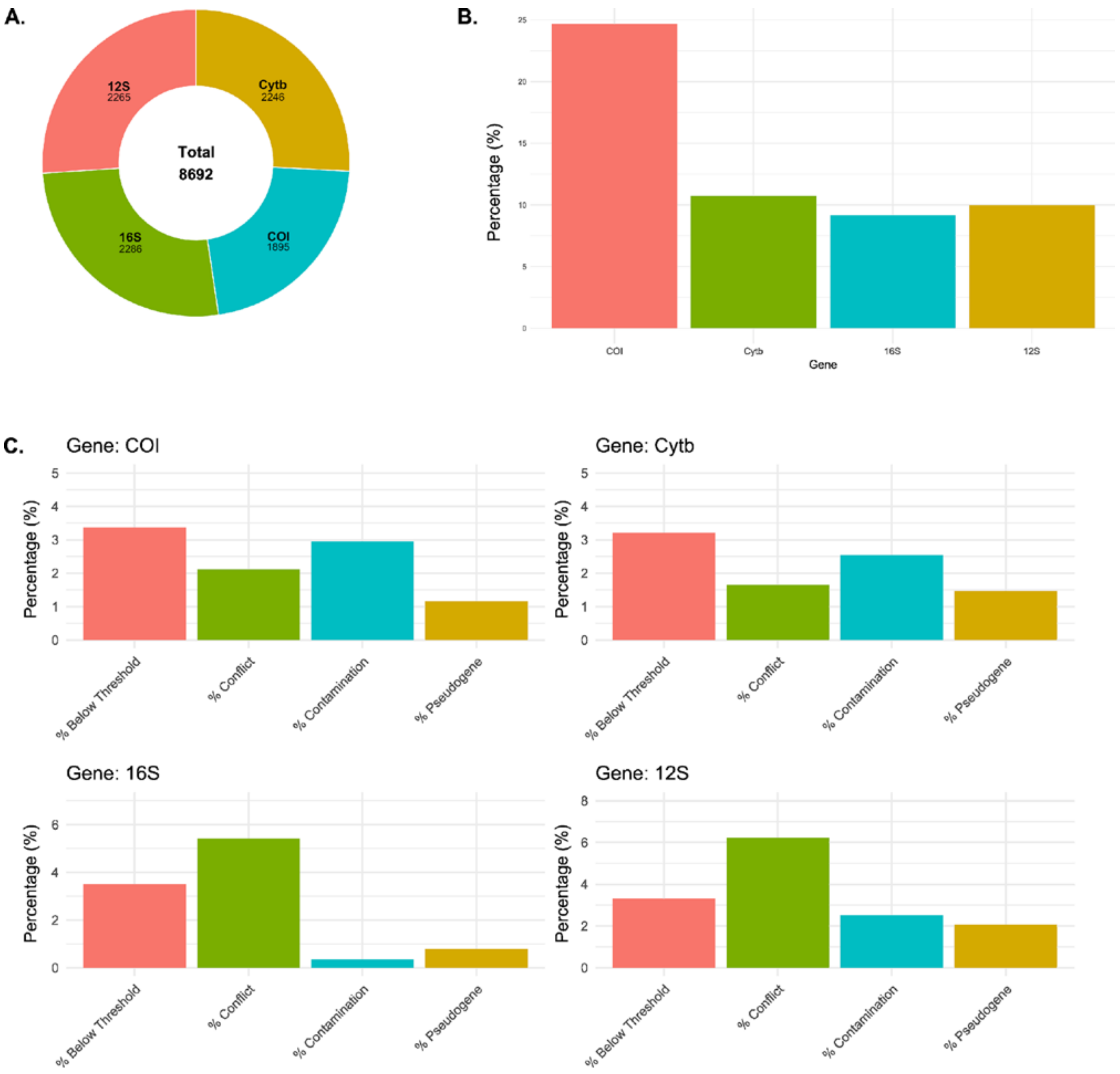


**Figure S29.** Amplification and sequencing patterns observed among the four mitochondrial genes used for DNA-typing bushmeat. A. Distribution of total nucleotide sequence numbers (N samples = 8692); B. Rates of PCR failure (lack of amplification); C. PCR biases. See the Methods section of the article for further context.

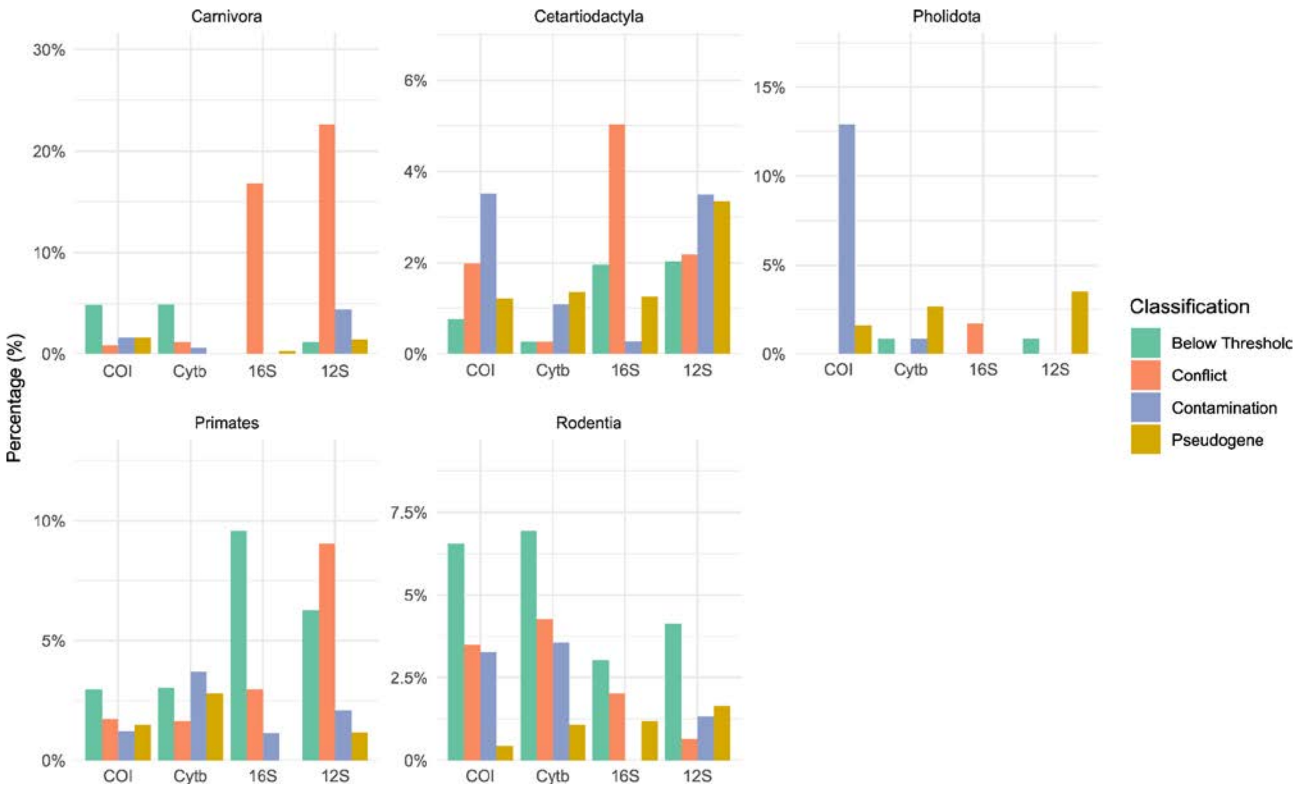


**Figure S30.** Amplification and sequencing patterns observed among the five main taxonomic orders among the four mitochondrial genes used for DNA-typing bushmeat. See Fig. S28 for extended legend.

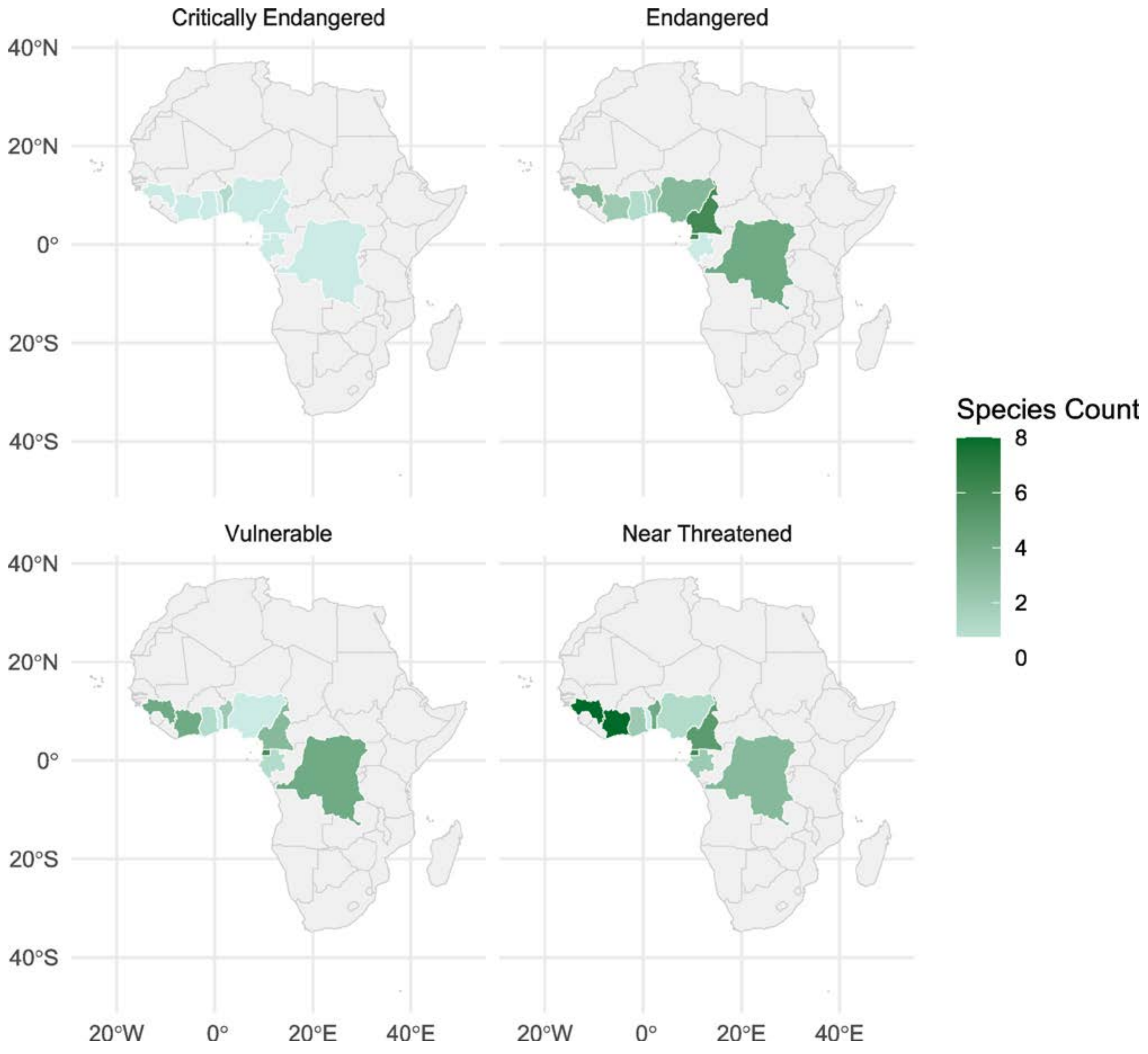


**Figure S31.** Distribution of species with a concerning conservation status in the IUCN Red List of threatened species across tropical African countries. Critically endangered (top left); Endangered (top right); Vulnerable (bottom left) and Neat Threatened (bottom right) species by IUCN conservation status across tropical African countries.

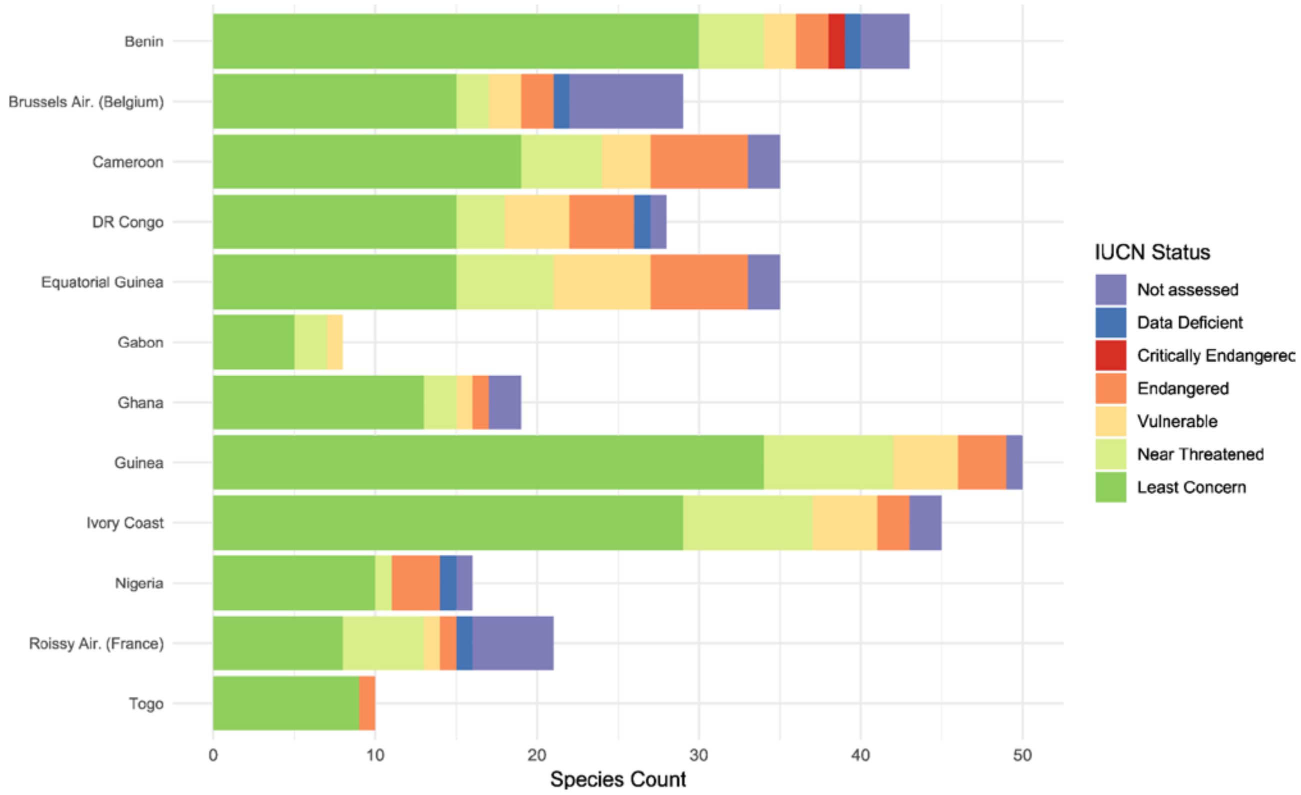


**Figure S32.** By country / airport distribution of IUCN Red List categories for bushmeat taxa.

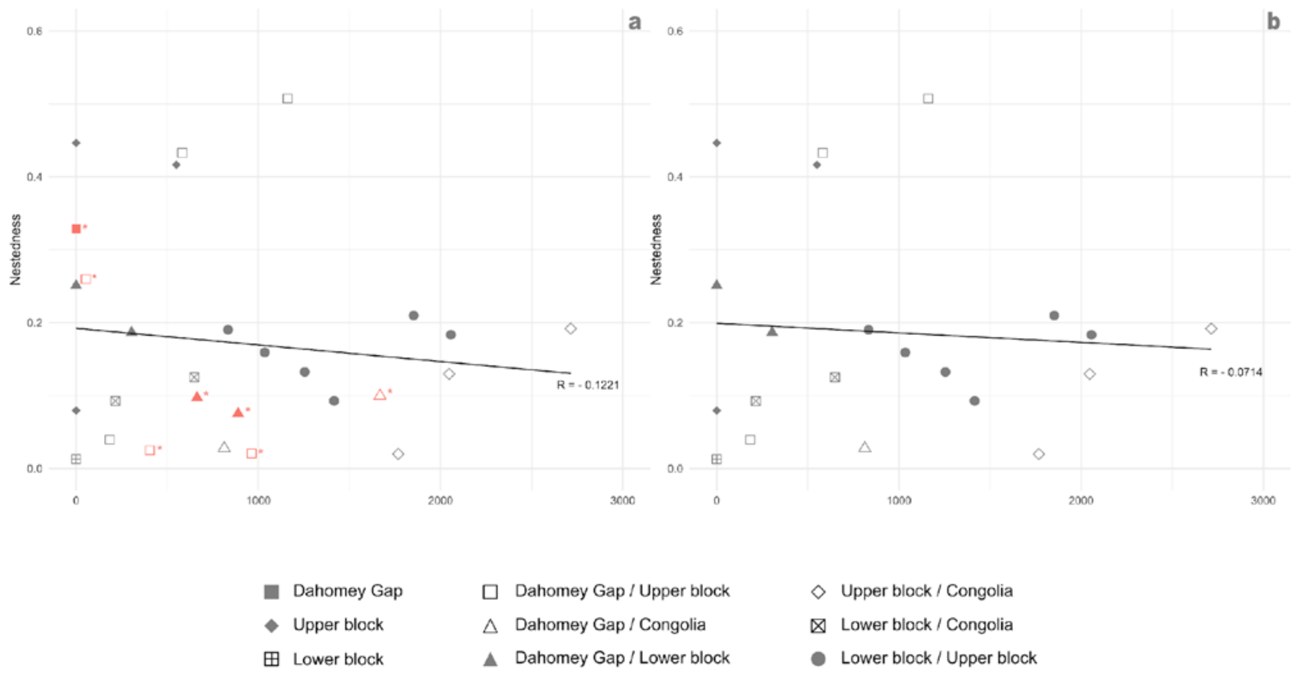


**Figure S33.** Correlation trends between phylogenetic nestedness (one of the components of phylogenetic β-diversity) and geographic distances as observed from national bushmeat markets. (a) including Benin; (b) excluding Benin.

**Table S1.** Metadata for the newly produced samples used in this study. [Excel table]

**Table S2.** Distribution of PCR failure rates across all genes and main taxonomic orders.

| **Taxonomic order** | **COI** | **Cytb** | **16S** | **12S** |
|---|---|---|---|---|
| **Carnivora** | 33.78% | 7.45% | 5.05% | 8.24% |
| **Cetartiodactyla** | 17.22% | 6.96% | 9.37% | 12.91% |
| **Pholidota** | 49.59% | 8.13% | 4.88% | 7.32% |
| **Primates** | 13.43% | 8.32% | 6.40% | 7.89% |
| **Rodentia** | 30.98% | 15.34% | 10.53% | 8.87% |
| **Average** | **29.00%** | **9.24%** | **7.24%** | **9.05%** |

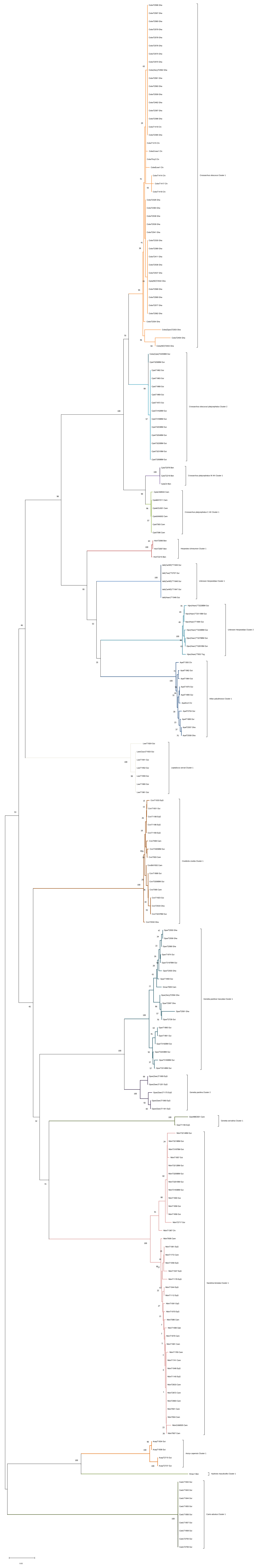
Crossarchus obscurus Cluster 1
Crossarchus obscurus/ platycephalus Cluster 2
Crossarchus platycephalus W Afr Cluster 1
Crossarchus platycephalus C Afr Cluster 1
Herpestes ichneumon Cluster 1
Unknown Herpestidae Cluster 1
Unknown Herpestidae Cluster 2
Atilax paludinosus Cluster 1
Leptailurus serval Cluster 1
Civettictis civetta Cluster 1
Genetta pardina/ maculata Cluster 1
Genetta pardina Cluster 2
Genetta servalina Cluster 1
Nandinia binotata Cluster 1
Aonyx capensis Cluster 1
Hydrictis maculicollis Cluster 1
Canis adustus Cluster 1

Figure_S1. Optimal COI Neighbour Joining tree of Carnivora bushmeat samples

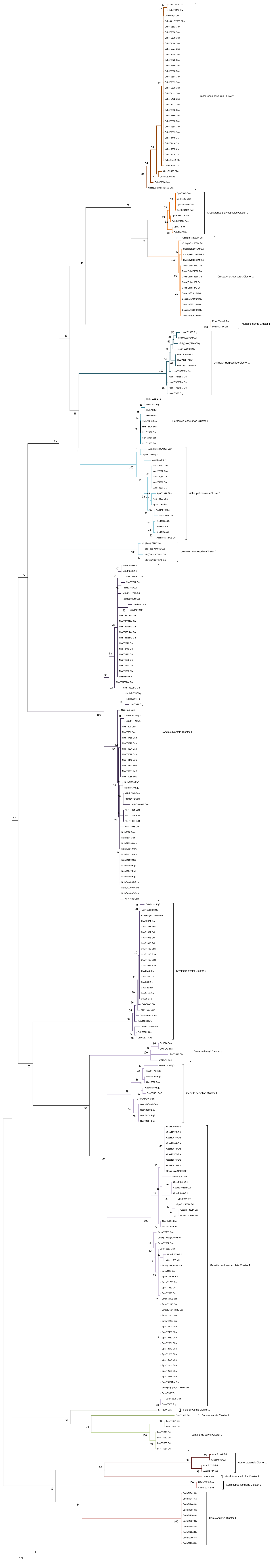

Crossarchus obscurus Cluster 1
Crossarchus platycephalus Cluster 1
Crossarchus obscurus Cluster 2
Mungos mungo Cluster 1
Unknown Herpestidae Cluster 1
Herpestes ichneumon Cluster 1
Atilax paludinosus Cluster 1
Unknown Herpestidae Cluster 2
Nandinia binotata Cluster 1
Civettictis civetta Cluster 1
Genetta thierryi Cluster 1
Genetta servalina Cluster 1
Genetta pardina/maculata Cluster 1
Felis silvestris Cluster 1
Caracal aurata Cluster 1
Leptailurus serval Cluster 1
Aonyx capensis Cluster 1
Hydrictis maculicollis Cluster 1
Canis lupus familiaris Cluster 1
Canis adustus Cluster 1
0.02

Figure_S2. Optimal COI Neighbour Joining tree of Carnivora bushmeat samples

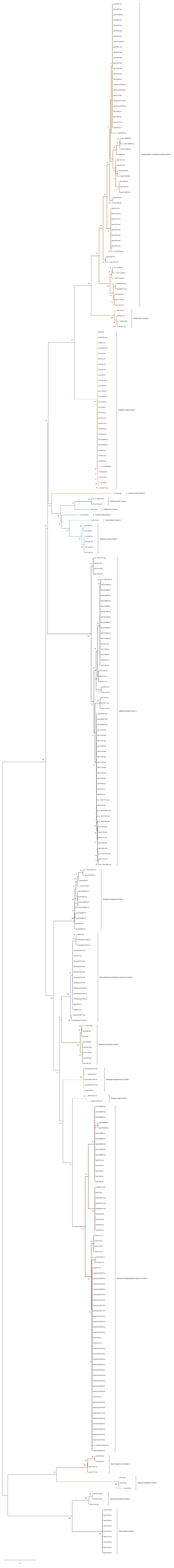

Figure_S3. Optimal 16S Neighbour Joining tree of Carnivora bushmeat samples.

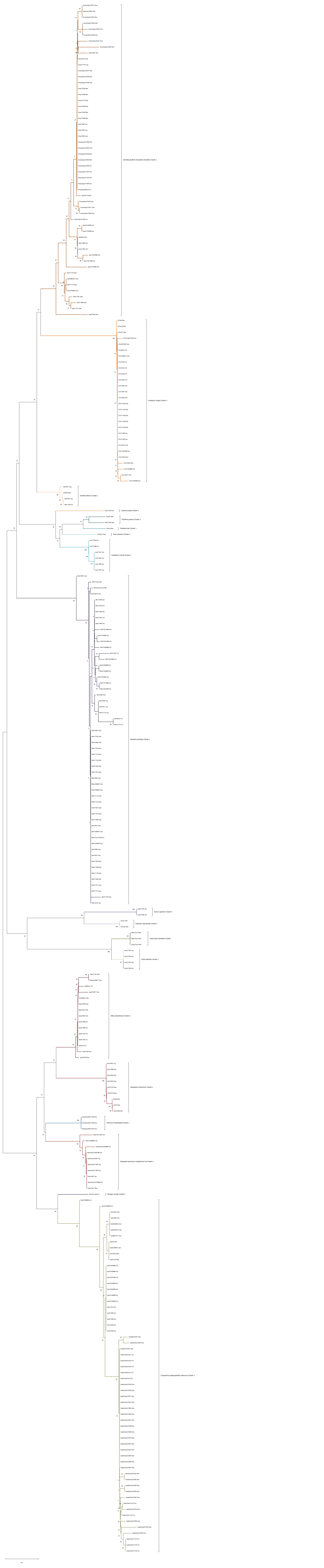

Genetta pardina/ maculata/ servalina Cluster 1
Civettictis civetta Cluster 1
Genetta thierryi Cluster 1
Caracal aurata Cluster 1
Panthera pardus Cluster 1
Panthera leo Cluster 1
Felis silvestris Cluster 1
Leptailurus serval Cluster 1
Nandinia binotata Cluster 1
Aonyx capensis Cluster 1
Hydrictis maculicollis Cluster 1
Canis lupus familiaris Cluster
Canis adustus Cluster 1
Atilax paludinosos Cluster 1
Herpestes ichneumon Cluster 1
Unknown Herpestidae Cluster 1
Herpestes javanicus/ sanguineus/ sp Cluster 1
Mungos mungo Cluster 1
Crossarchus platycephalus/ obscurus Cluster 1

Figure_S4. Optimal 12S Neighbour Joining tree of Carnivora bushmeat samples.

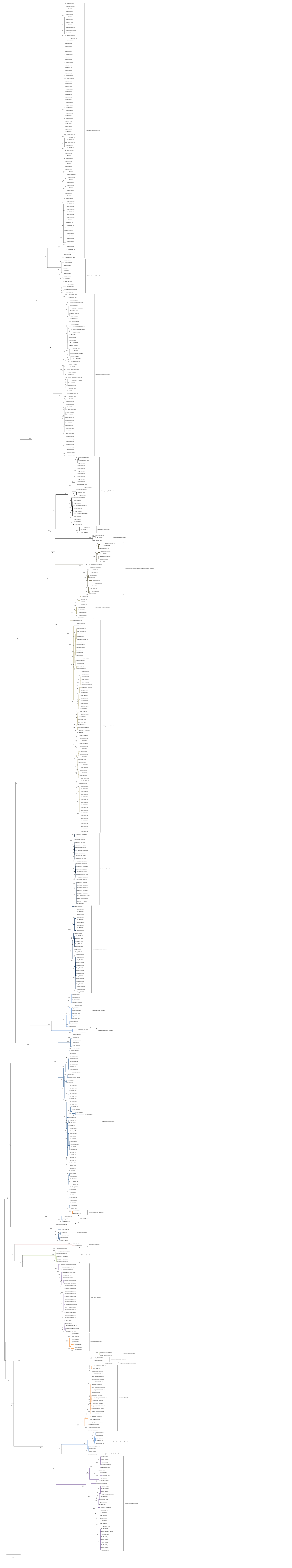

Figure_S5. Optimal COI Neighbour Joining tree of Cetartiodactyla bushmeat samples.

Figure_S6. Optimal Cytb Neighbour Joining tree of Cetartiodactyla bushmeat samples.

Figure_S7. Optimal 16S Neighbour Joining tree of Cetartiodactyla bushmeat samples.

Figure_S8. Optimal 12S Neighbour Joining tree of Cetartiodactyla bushmeat samples.

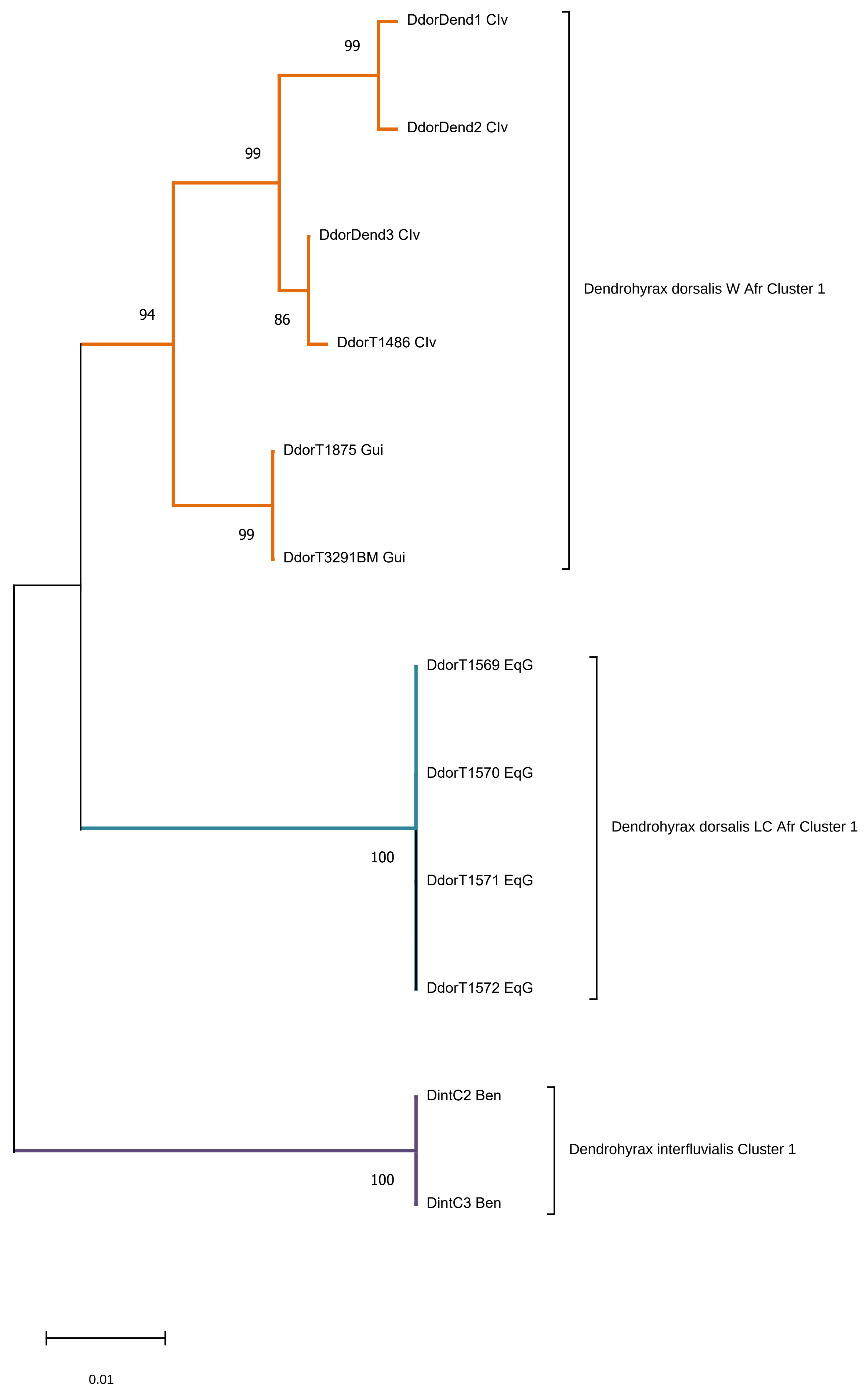
DdorDend1 CIv
99
DdorDend2 CIv
99
DdorDend3 CIv
94
86
DdorT1486 CIv
DdorT1875 Gui
99
DdorT3291BM Gui
Dendrohyrax dorsalis W Afr Cluster 1
DdorT1569 EqG
DdorT1570 EqG
100
DdorT1571 EqG
DdorT1572 EqG
Dendrohyrax dorsalis LC Afr Cluster 1
DintC2 Ben
100
DintC3 Ben
Dendrohyrax interfluvialis Cluster 1
0.01

Figure_S9. Optimal COI Neighbour Joining tree of Hyracoidea bushmeat samples.

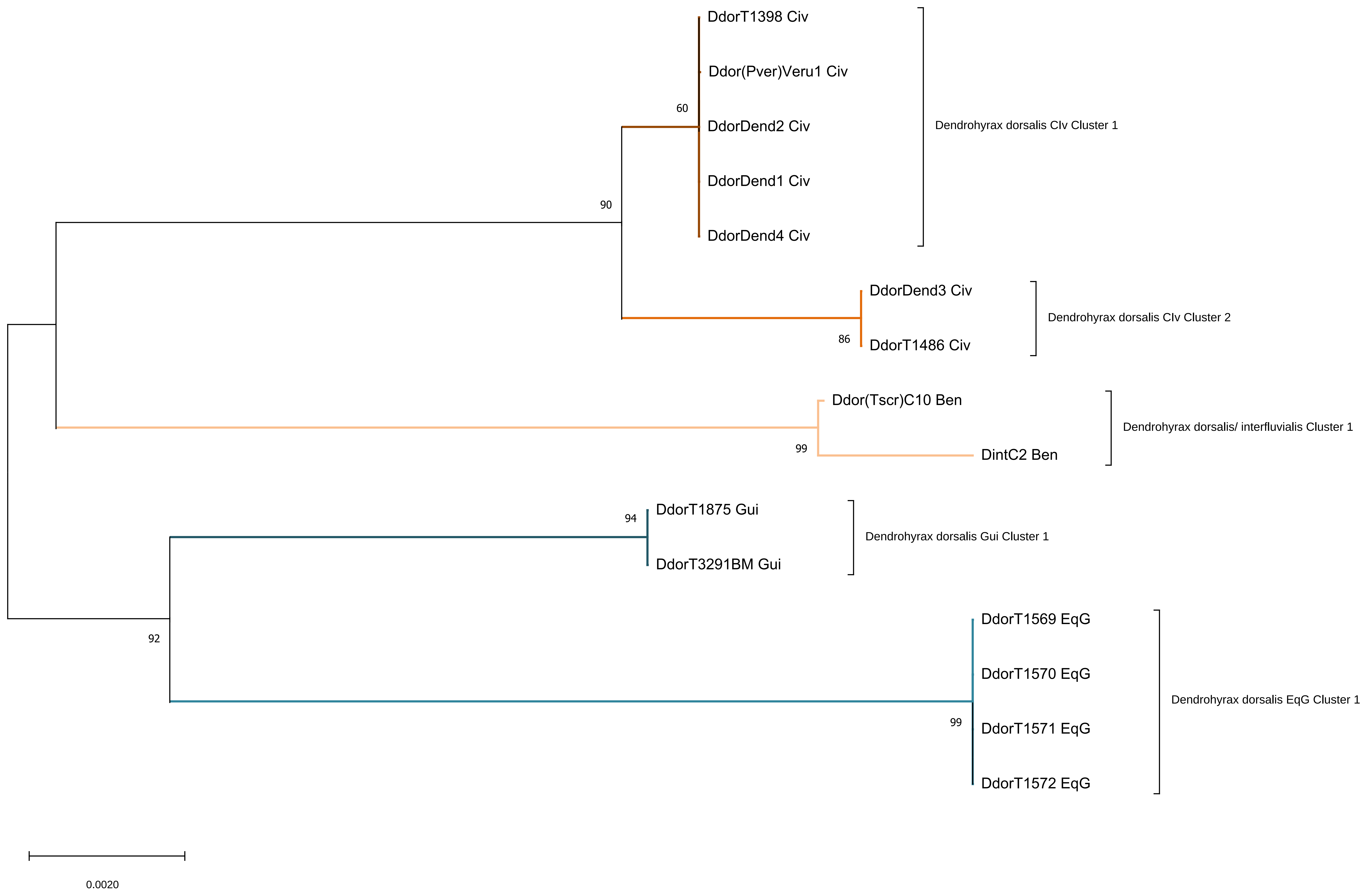
DdorT1398 Civ
Ddor(Pver)Veru1 Civ
60
DdorDend2 Civ
DdorDend1 Civ
90
DdorDend4 Civ
Dendrohyrax dorsalis CIv Cluster 1
DdorDend3 Civ
86
DdorT1486 Civ
Dendrohyrax dorsalis CIv Cluster 2
Ddor(Tscr)C10 Ben
99
DintC2 Ben
Dendrohyrax dorsalis/ interfluvialis Cluster 1
94
DdorT1875 Gui
DdorT3291BM Gui
Dendrohyrax dorsalis Gui Cluster 1
92
DdorT1569 EqG
DdorT1570 EqG
99
DdorT1571 EqG
DdorT1572 EqG
Dendrohyrax dorsalis EqG Cluster 1
0.0020

Figure_S10. Optimal 16S Neighbour Joining tree of Hyracoidea bushmeat samples.

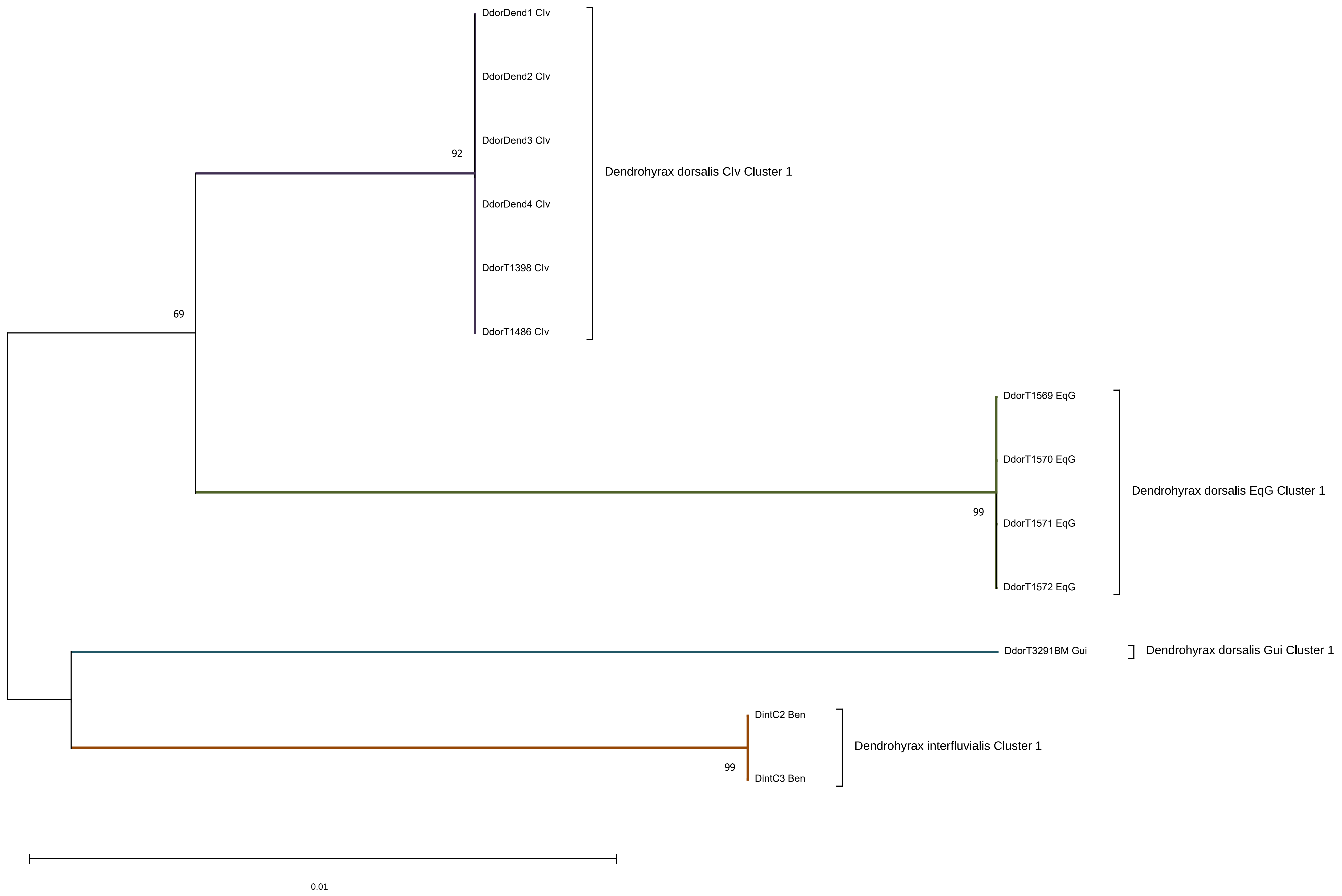
DdorDend1 CIv
DdorDend2 CIv
DdorDend3 CIv
DdorDend4 CIv
DdorT1398 CIv
DdorT1486 CIv
92
69
Dendrohyrax dorsalis CIv Cluster 1
DdorT1569 EqG
DdorT1570 EqG
DdorT1571 EqG
DdorT1572 EqG
99
Dendrohyrax dorsalis EqG Cluster 1
DdorT3291BM Gui
Dendrohyrax dorsalis Gui Cluster 1
DintC2 Ben
DintC3 Ben
99
Dendrohyrax interfluvialis Cluster 1
0.01

Figure_S11. Optimal 12S Neighbour Joining tree of Hyracoidea bushmeat samples.

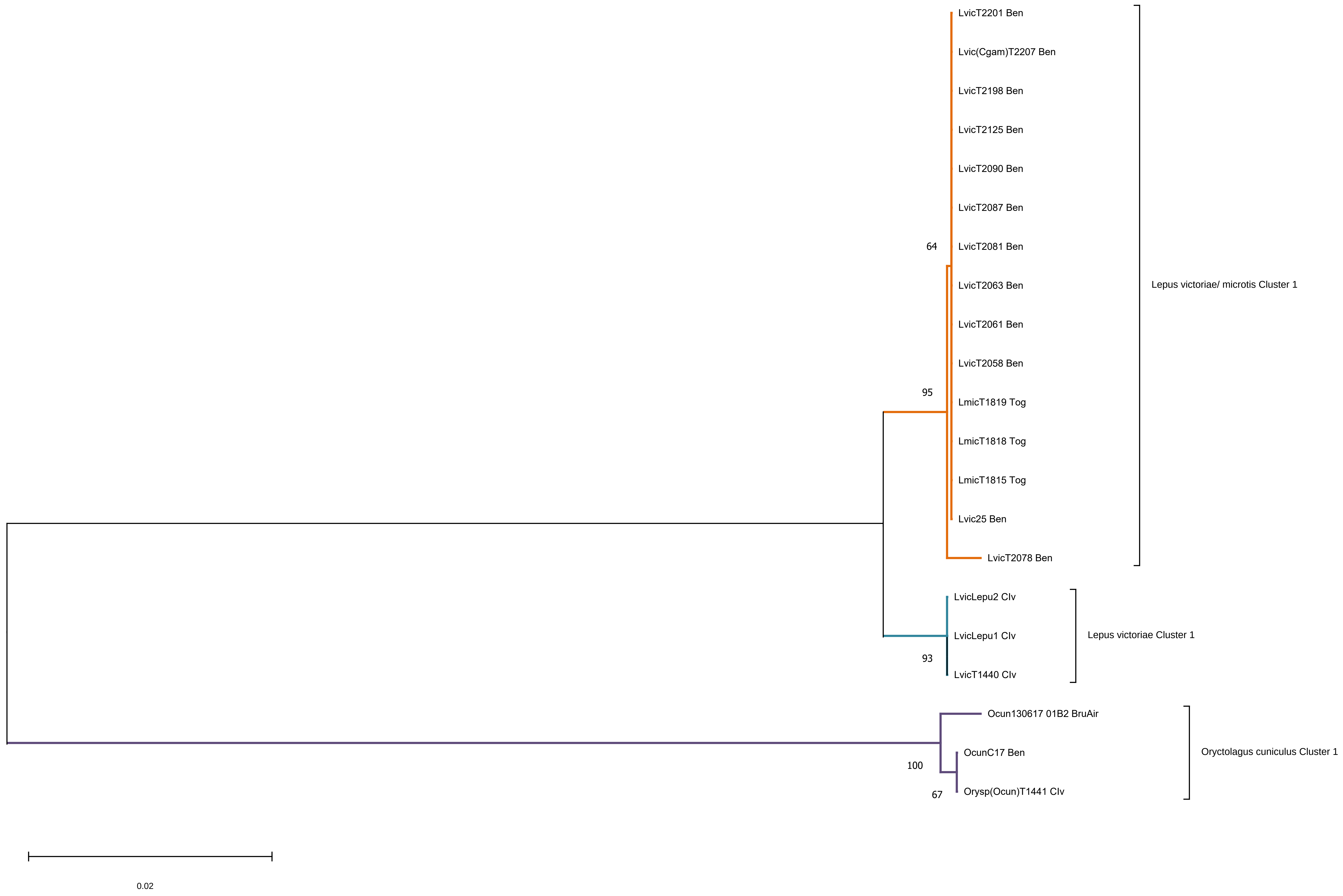
LvicT2201 Ben
Lvic(Cgam)T2207 Ben
LvicT2198 Ben
LvicT2125 Ben
LvicT2090 Ben
LvicT2087 Ben
64
LvicT2081 Ben
LvicT2063 Ben
LvicT2061 Ben
LvicT2058 Ben
95
LmicT1819 Tog
LmicT1818 Tog
LmicT1815 Tog
Lvic25 Ben
LvicT2078 Ben
Lepus victoriae/ microtis Cluster 1
LvicLepu2 Clv
LvicLepu1 Clv
93
LvicT1440 Clv
Lepus victoriae Cluster 1
Ocun130617 01B2 BruAir
OcunC17 Ben
100
67
Orysp(Ocun)T1441 Clv
Oryctolagus cuniculus Cluster 1
0.02

Figure_S12. Optimal COI Neighbour Joining tree of Lagomorpha bushmeat samples.

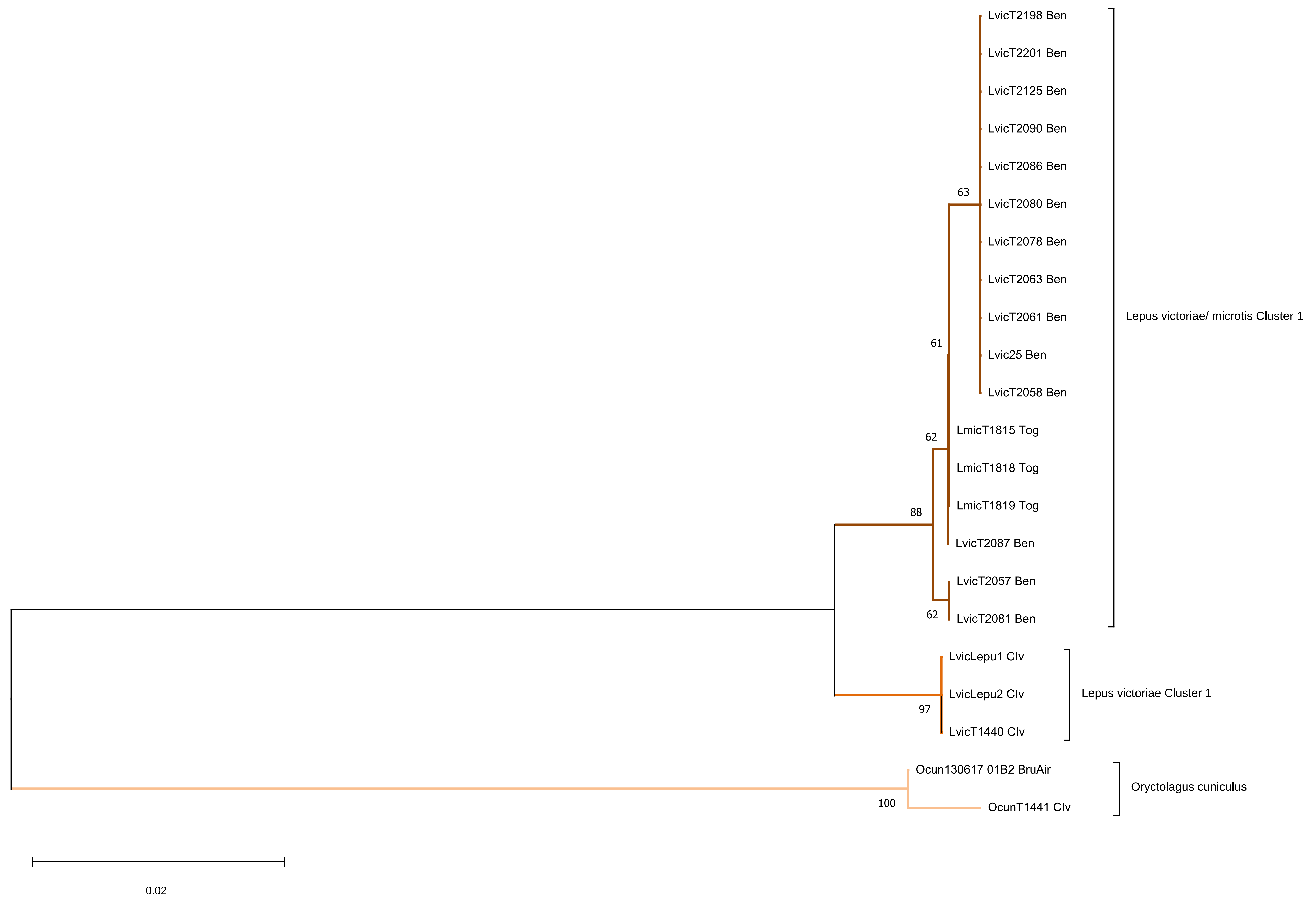
LvicT2198 Ben
LvicT2201 Ben
LvicT2125 Ben
LvicT2090 Ben
LvicT2086 Ben
63
LvicT2080 Ben
LvicT2078 Ben
LvicT2063 Ben
LvicT2061 Ben
Lepus victoriae/ microtis Cluster 1
61
Lvic25 Ben
LvicT2058 Ben
LmicT1815 Tog
62
LmicT1818 Tog
88
LmicT1819 Tog
LvicT2087 Ben
LvicT2057 Ben
62
LvicT2081 Ben
LvicLepu1 Clv
LvicLepu2 Clv
Lepus victoriae Cluster 1
97
LvicT1440 Clv
Ocun130617 01B2 BruAir
Oryctolagus cuniculus
100
OcunT1441 Clv
0.02

Figure_S13. Optimal Cytb Neighbour Joining tree of Lagomorpha bushmeat samples.

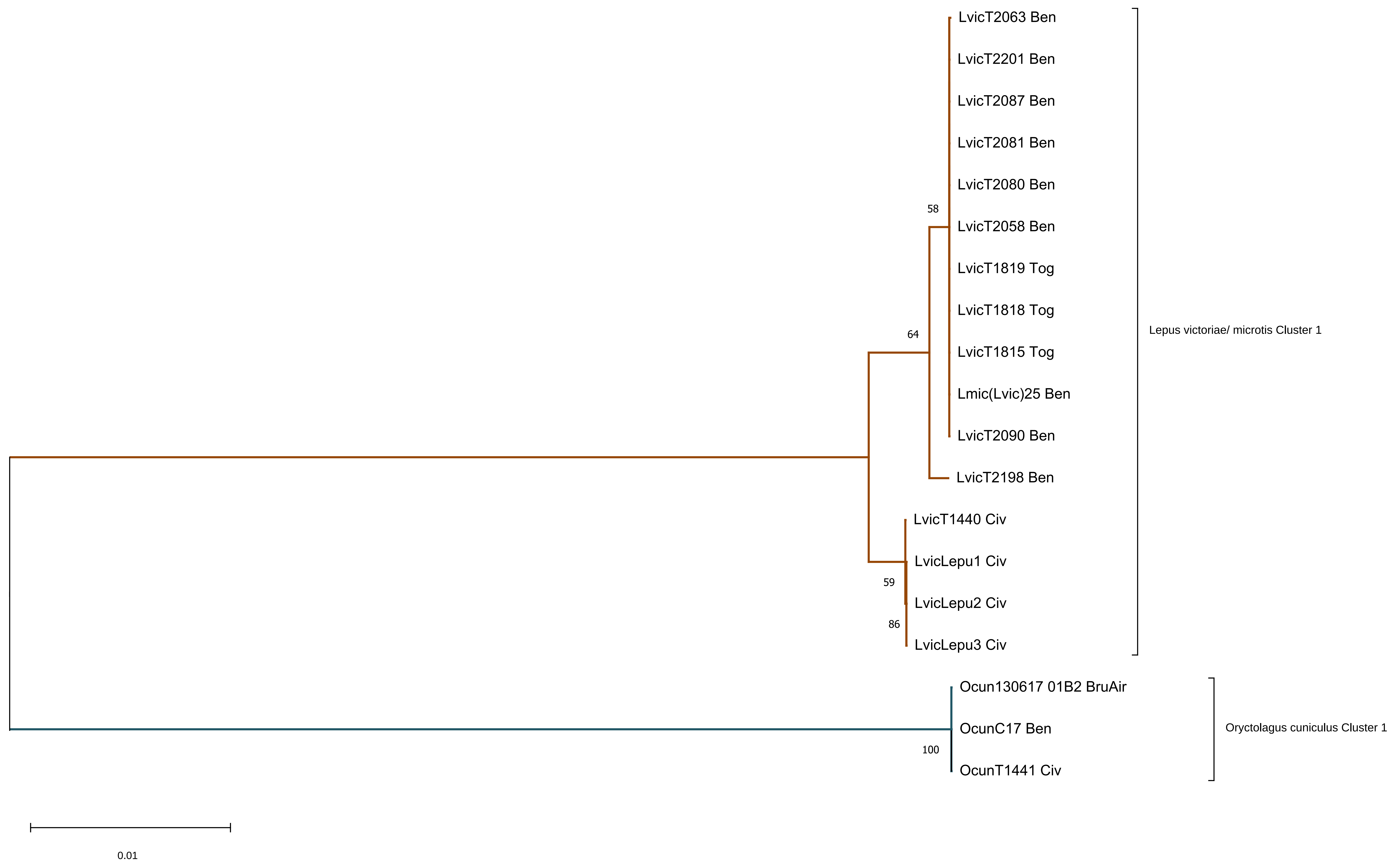

LvicT2063 Ben
LvicT2201 Ben
LvicT2087 Ben
LvicT2081 Ben
LvicT2080 Ben
58
LvicT2058 Ben
LvicT1819 Tog
LvicT1818 Tog
64
LvicT1815 Tog
Lmic(Lvic)25 Ben
LvicT2090 Ben
LvicT2198 Ben
LvicT1440 Civ
LvicLepu1 Civ
59
LvicLepu2 Civ
86
LvicLepu3 Civ
Lepus victoriae/ microtis Cluster 1
Ocun130617 01B2 BruAir
OcunC17 Ben
100
OcunT1441 Civ
Oryctolagus cuniculus Cluster 1
0.01

Figure_S14. Optimal 16S Neighbour Joining tree of Lagomorpha bushmeat samples.

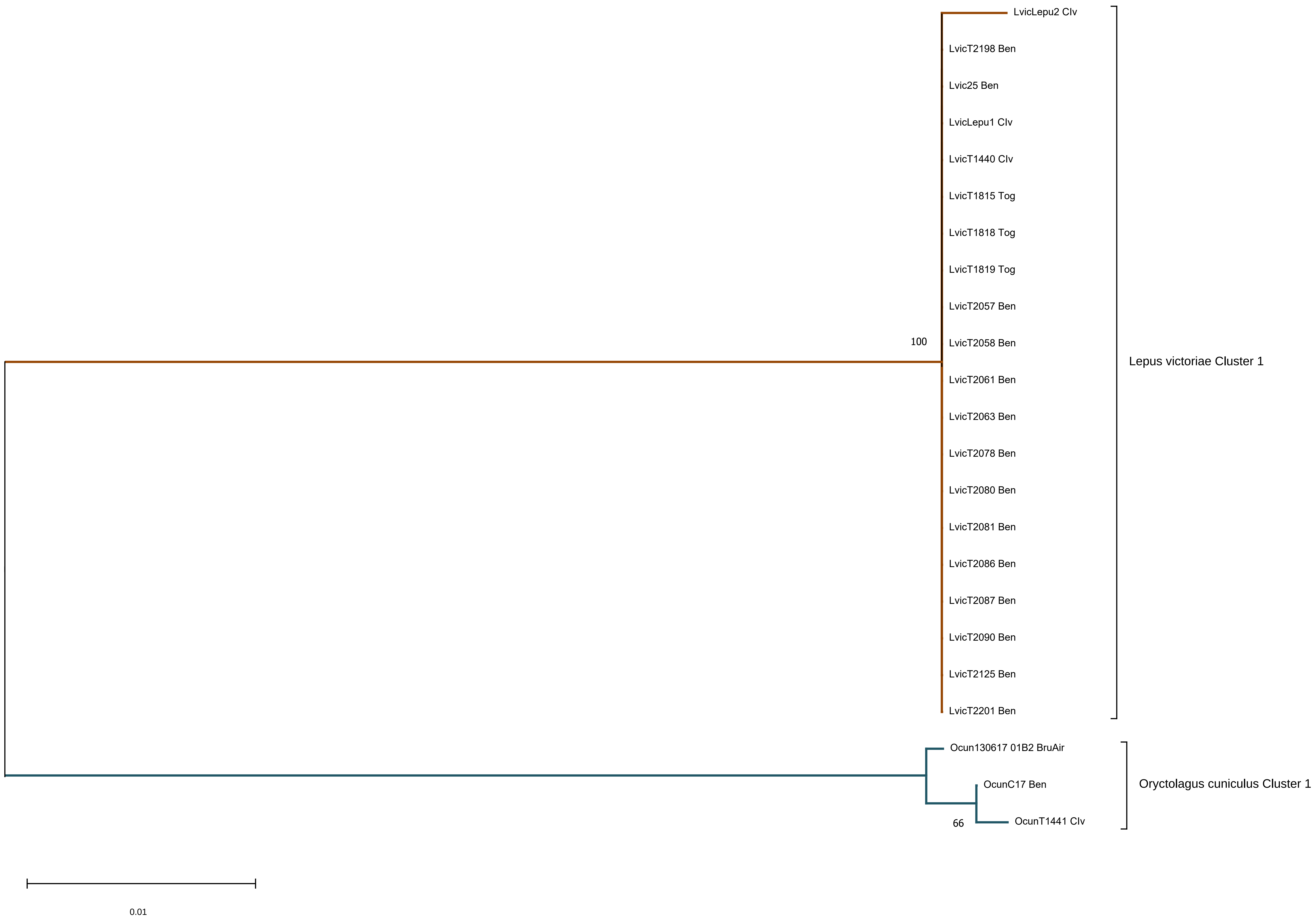
LvicLepu2 Clv
LvicT2198 Ben
Lvic25 Ben
LvicLepu1 Clv
LvicT1440 Clv
LvicT1815 Tog
LvicT1818 Tog
LvicT1819 Tog
LvicT2057 Ben
100
LvicT2058 Ben
LvicT2061 Ben
LvicT2063 Ben
LvicT2078 Ben
LvicT2080 Ben
LvicT2081 Ben
LvicT2086 Ben
LvicT2087 Ben
LvicT2090 Ben
LvicT2125 Ben
LvicT2201 Ben
Lepus victoriae Cluster 1
Ocun130617 01B2 BruAir
OcunC17 Ben
66
OcunT1441 Clv
Oryctolagus cuniculus Cluster 1
0.01

Figure_S15. Optimal 12S Neighbour Joining tree of Lagomorpha bushmeat samples.

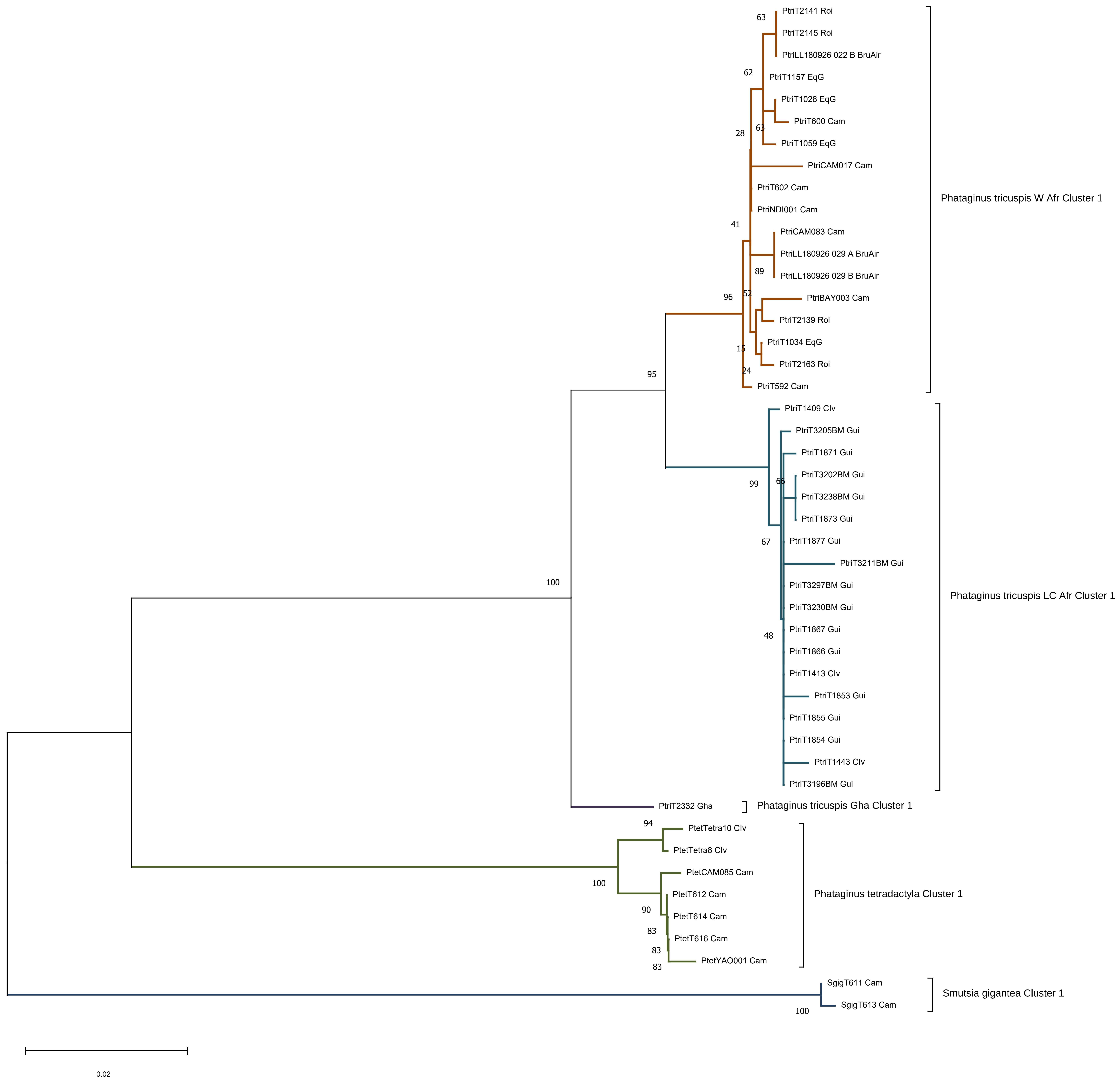
PtriT2141 Roi
PtriT2145 Roi
PtriLL180926 022 B BruAir
PtriT1157 EqG
PtriT1028 EqG
PtriT600 Cam
PtriT1059 EqG
PtriCAM017 Cam
PtriT602 Cam
PtriNDI001 Cam
PtriCAM083 Cam
PtriLL180926 029 A BruAir
PtriLL180926 029 B BruAir
PtriBAY003 Cam
PtriT2139 Roi
PtriT1034 EqG
PtriT2163 Roi
PtriT592 Cam
Phataginus tricuspis W Afr Cluster 1
PtriT1409 Clv
PtriT3205BM Gui
PtriT1871 Gui
PtriT3202BM Gui
PtriT3238BM Gui
PtriT1873 Gui
PtriT1877 Gui
PtriT3211BM Gui
PtriT3297BM Gui
PtriT3230BM Gui
PtriT1867 Gui
PtriT1866 Gui
PtriT1413 Clv
PtriT1853 Gui
PtriT1855 Gui
PtriT1854 Gui
PtriT1443 Clv
PtriT3196BM Gui
Phataginus tricuspis LC Afr Cluster 1
PtriT2332 Gha
Phataginus tricuspis Gha Cluster 1
PtetTetra10 Clv
PtetTetra8 Clv
PtetCAM085 Cam
PtetT612 Cam
PtetT614 Cam
PtetT616 Cam
PtetYAO001 Cam
Phataginus tetradactyla Cluster 1
SgigT611 Cam
SgigT613 Cam
Smutsia gigantea Cluster 1
63
62
63
28
41
89
96
52
15
24
95
99
66
67
48
100
94
100
90
83
83
83
100
0.02

Figure_S16. Optimal COI Neighbour Joining tree of Pholidota bushmeat samples.

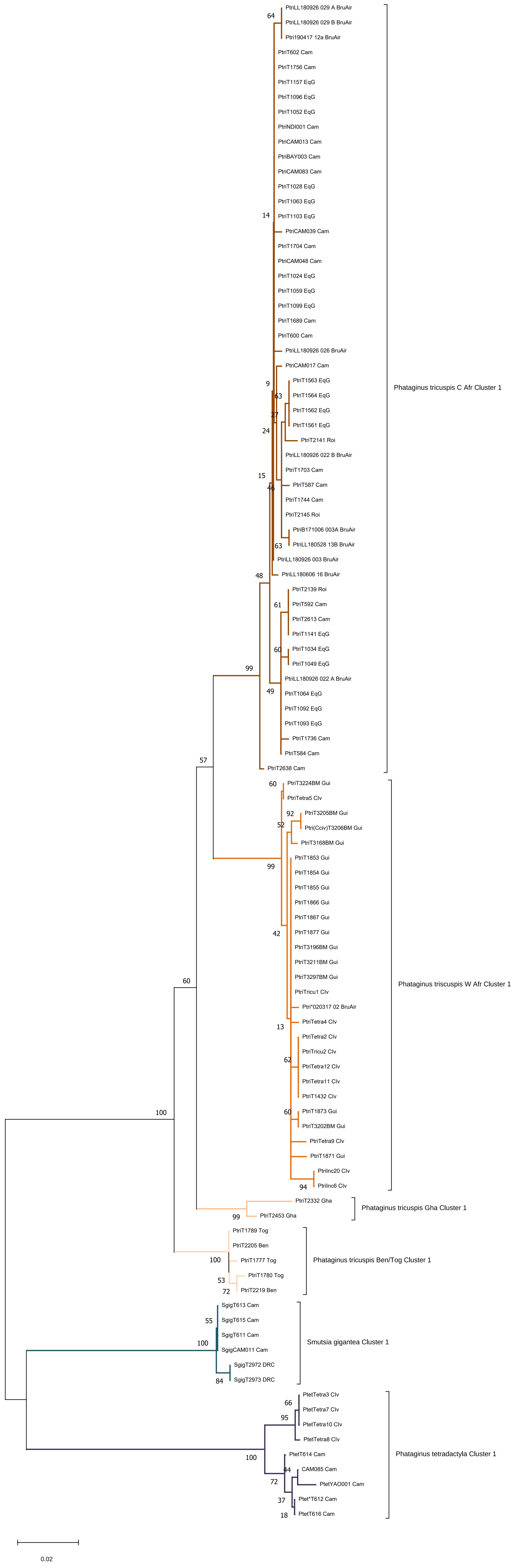
PtriLL180926 029 A BruAir
PtriLL180926 029 B BruAir
Ptri190417 12a BruAir
PtriT602 Cam
PtriT1756 Cam
PtriT1157 EqG
PtriT1096 EqG
PtriT1052 EqG
PtriNDI001 Cam
PtriCAM013 Cam
PtriBAY003 Cam
PtriCAM083 Cam
PtriT1028 EqG
PtriT1063 EqG
PtriT1103 EqG
PtriCAM039 Cam
PtriT1704 Cam
PtriCAM048 Cam
PtriT1024 EqG
PtriT1059 EqG
PtriT1099 EqG
PtriT1689 Cam
PtriT600 Cam
PtriLL180926 026 BruAir
PtriCAM017 Cam
PtriT1563 EqG
PtriT1564 EqG
PtriT1562 EqG
PtriT1561 EqG
PtriT2141 Roi
PtriLL180926 022 B BruAir
PtriT1703 Cam
PtriT587 Cam
PtriT1744 Cam
PtriT2145 Roi
PtriB171006 003A BruAir
PtriLL180528 13B BruAir
PtriLL180926 003 BruAir
PtriLL180606 16 BruAir
PtriT2139 Roi
PtriT592 Cam
PtriT2613 Cam
PtriT1141 EqG
PtriT1034 EqG
PtriT1049 EqG
PtriLL180926 022 A BruAir
PtriT1064 EqG
PtriT1092 EqG
PtriT1093 EqG
PtriT1736 Cam
PtriT584 Cam
PtriT2638 Cam
Phataginus tricuspis C Afr Cluster 1
PtriT3224BM Gui
PtriTetra5 CIv
PtriT3205BM Gui
Ptri(Cciv)T3206BM Gui
PtriT3168BM Gui
PtriT1853 Gui
PtriT1854 Gui
PtriT1855 Gui
PtriT1866 Gui
PtriT1867 Gui
PtriT1877 Gui
PtriT3196BM Gui
PtriT3211BM Gui
PtriT3297BM Gui
PtriTricu1 CIv
Ptri*020317 02 BruAir
PtriTetra4 CIv
PtriTetra2 CIv
PtriTricu2 CIv
PtriTetra12 CIv
PtriTetra11 CIv
PtriT1432 CIv
PtriT1873 Gui
PtriT3202BM Gui
PtriTetra9 CIv
PtriT1871 Gui
PtriInc20 CIv
PtriInc6 CIv
Phataginus triscuspis W Afr Cluster 1
PtriT2332 Gha
PtriT2453 Gha
Phataginus tricuspis Gha Cluster 1
PtriT1789 Tog
PtriT2205 Ben
PtriT1777 Tog
PtriT1780 Tog
PtriT2219 Ben
Phataginus tricuspis Ben/Tog Cluster 1
SgigT613 Cam
SgigT615 Cam
SgigT611 Cam
SgigCAM011 Cam
SgigT2972 DRC
SgigT2973 DRC
Smutsia gigantea Cluster 1
PtetTetra3 CIv
PtetTetra7 CIv
PtetTetra10 CIv
PtetTetra8 CIv
PtetT614 Cam
CAM085 Cam
PtetYAO001 Cam
Ptet*T612 Cam
PtetT616 Cam
Phataginus tetradactyla Cluster 1
0.02

Figure_S17. Optimal Cytb Neighbour Joining tree of Pholidota bushmeat samples.

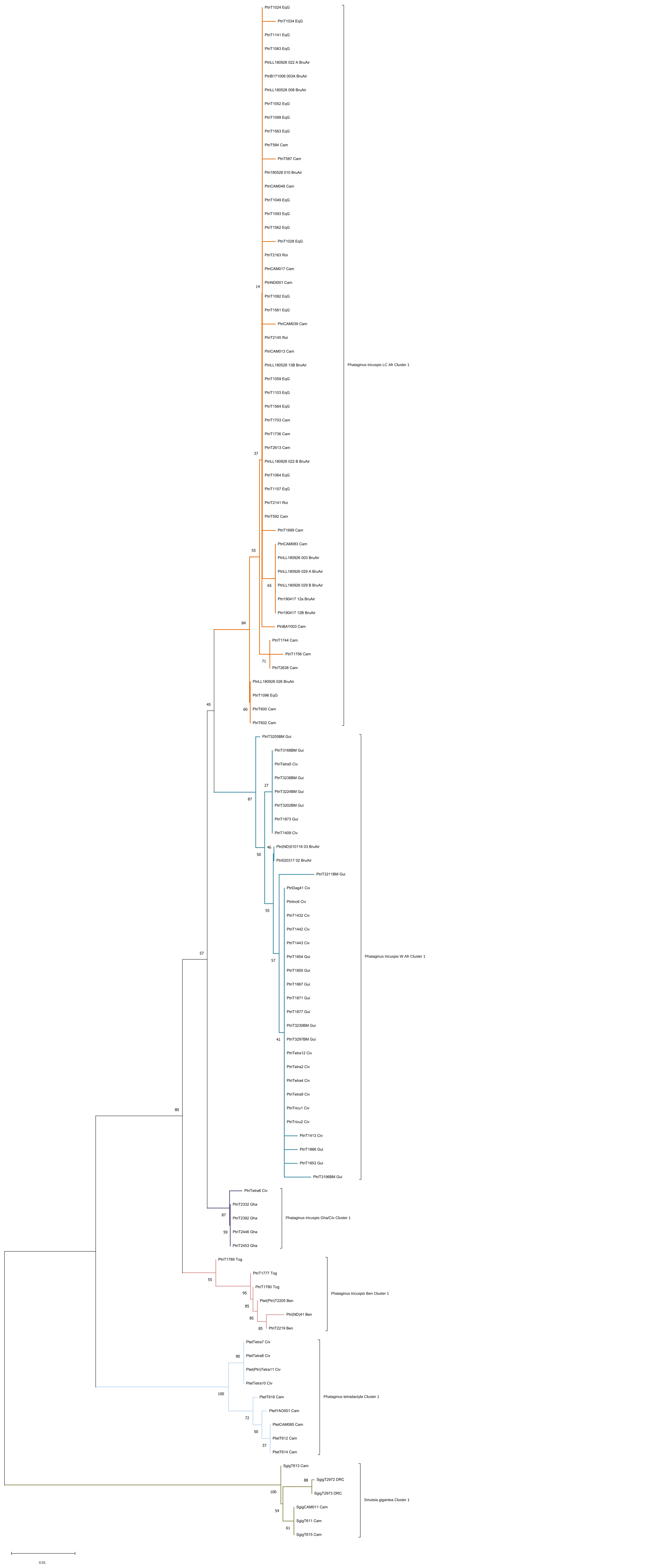

PtriT1024 EqG
PtriT1034 EqG
PtriT1141 EqG
PtriT1063 EqG
PtriLL180926 022 A BruAir
PtriB171006 003A BruAir
PtriLL180528 008 BruAir
PtriT1052 EqG
PtriT1099 EqG
PtriT1563 EqG
PtriT584 Cam
PtriT587 Cam
Ptri180528 010 BruAir
PtriCAM048 Cam
PtriT1049 EqG
PtriT1093 EqG
PtriT1562 EqG
PtriT1028 EqG
PtriT2163 Roi
PtriCAM017 Cam
PtriNDI001 Cam
PtriT1092 EqG
PtriT1561 EqG
PtriCAM039 Cam
PtriT2145 Roi
PtriCAM013 Cam
PtriLL180528 13B BruAir
PtriT1059 EqG
PtriT1103 EqG
PtriT1564 EqG
PtriT1703 Cam
PtriT1736 Cam
PtriT2613 Cam
PtriLL180926 022 B BruAir
PtriT1064 EqG
PtriT1157 EqG
PtriT2141 Roi
PtriT592 Cam
PtriT1689 Cam
PtriCAM083 Cam
PtriLL180926 003 BruAir
PtriLL180926 029 A BruAir
PtriLL180926 029 B BruAir
Ptri190417 12a BruAir
Ptri190417 12B BruAir
PtriBAY003 Cam
PtriT1744 Cam
PtriT1756 Cam
PtriT2638 Cam
PtriLL180926 026 BruAir
PtriT1096 EqG
PtriT600 Cam
PtriT602 Cam
Phataginus tricuspis LC Afr Cluster 1
PtriT3205BM Gui
PtriT3168BM Gui
PtriTetra5 Civ
PtriT3238BM Gui
PtriT3224BM Gui
PtriT3202BM Gui
PtriT1873 Gui
PtriT1409 Civ
Ptri(ND)010116 03 BruAir
Ptri020317 02 BruAir
PtriT3211BM Gui
PtriDag41 Civ
PtriInc6 Civ
PtriT1432 Civ
PtriT1442 Civ
PtriT1443 Civ
PtriT1854 Gui
PtriT1855 Gui
PtriT1867 Gui
PtriT1871 Gui
PtriT1877 Gui
PtriT3230BM Gui
PtriT3297BM Gui
PtriTetra12 Civ
PtriTetra2 Civ
PtriTetra4 Civ
PtriTetra9 Civ
PtriTricu1 Civ
PtriTricu2 Civ
PtriT1413 Civ
PtriT1866 Gui
PtriT1853 Gui
PtriT3196BM Gui
Phataginus tricuspis W Afr Cluster 1
PtriTetra6 Civ
PtriT2332 Gha
PtriT2392 Gha
PtriT2446 Gha
PtriT2453 Gha
Phataginus tricuspis Gha/Civ Cluster 1
PtriT1789 Tog
PtriT1777 Tog
PtriT1780 Tog
Ptet(Ptri)T2205 Ben
Ptri(ND)41 Ben
PtriT2219 Ben
Phataginus tricuspis Ben Cluster 1
PtetTetra7 Civ
PtetTetra8 Civ
Ptet(Ptri)Tetra11 Civ
PtetTetra10 Civ
PtetT616 Cam
PtetYAO001 Cam
PtetCAM085 Cam
PtetT612 Cam
PtetT614 Cam
Phataginus tetradactyla Cluster 1
SgigT613 Cam
SgigT2972 DRC
SgigT2973 DRC
SgigCAM011 Cam
SgigT611 Cam
SgigT615 Cam
Smutsia gigantea Cluster 1
14
37
55
65
84
71
60
45
87
27
46
50
55
57
41
57
80
87
59
55
95
85
85
85
90
100
72
50
37
88
100
54
61
0.01

Figure_S18. Optimal 16S Neighbour Joining tree of Pholidota bushmeat samples.

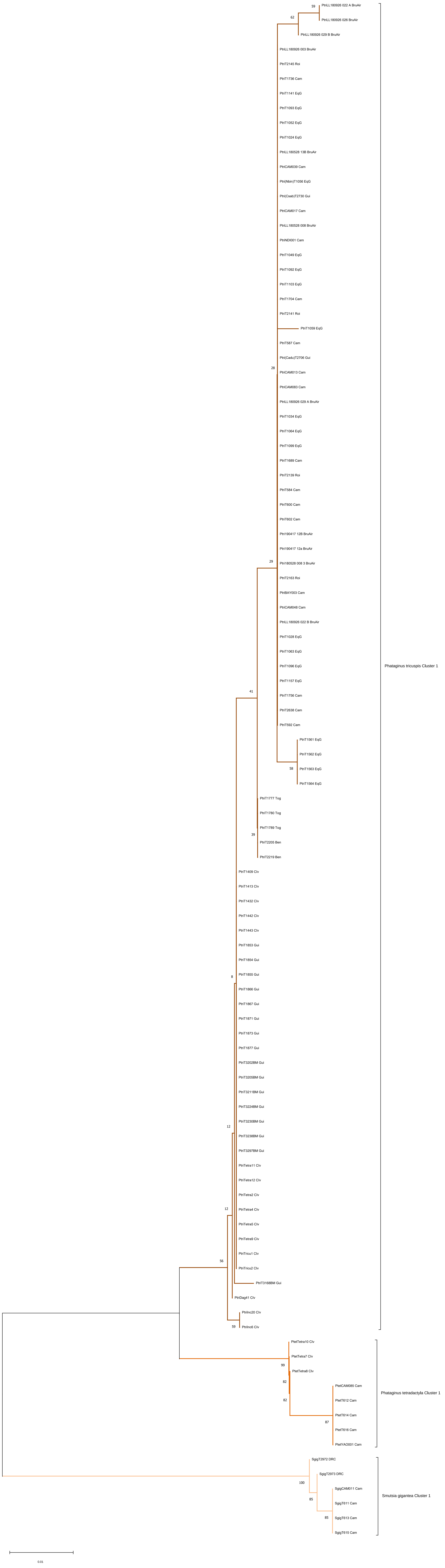
PtriLL180926 022 A BruAir
PtriLL180926 026 BruAir
PtriLL180926 029 B BruAir
PtriLL180926 003 BruAir
PtriT2145 Roi
PtriT1736 Cam
PtriT1141 EqG
PtriT1093 EqG
PtriT1052 EqG
PtriT1024 EqG
PtriLL180528 13B BruAir
PtriCAM039 Cam
Ptri(Nbin)T1056 EqG
Ptri(Csab)T2730 Gui
PtriCAM017 Cam
PtriLL180528 008 BruAir
PtriNDI001 Cam
PtriT1049 EqG
PtriT1092 EqG
PtriT1103 EqG
PtriT1704 Cam
PtriT2141 Roi
PtriT1059 EqG
PtriT587 Cam
Ptri(Cadu)T2706 Gui
PtriCAM013 Cam
PtriCAM083 Cam
PtriLL180926 029 A BruAir
PtriT1034 EqG
PtriT1064 EqG
PtriT1099 EqG
PtriT1689 Cam
PtriT2139 Roi
PtriT584 Cam
PtriT600 Cam
PtriT602 Cam
Ptri190417 12B BruAir
Ptri190417 12a BruAir
Ptri180528 008 3 BruAir
PtriT2163 Roi
PtriBAY003 Cam
PtriCAM048 Cam
PtriLL180926 022 B BruAir
PtriT1028 EqG
PtriT1063 EqG
PtriT1096 EqG
PtriT1157 EqG
PtriT1756 Cam
PtriT2638 Cam
PtriT592 Cam
PtriT1561 EqG
PtriT1562 EqG
PtriT1563 EqG
PtriT1564 EqG
PtriT1777 Tog
PtriT1780 Tog
PtriT1789 Tog
PtriT2205 Ben
PtriT2219 Ben
PtriT1409 Civ
PtriT1413 Civ
PtriT1432 Civ
PtriT1442 Civ
PtriT1443 Civ
PtriT1853 Gui
PtriT1854 Gui
PtriT1855 Gui
PtriT1866 Gui
PtriT1867 Gui
PtriT1871 Gui
PtriT1873 Gui
PtriT1877 Gui
PtriT3202BM Gui
PtriT3205BM Gui
PtriT3211BM Gui
PtriT3224BM Gui
PtriT3230BM Gui
PtriT3238BM Gui
PtriT3297BM Gui
PtriTetra11 Civ
PtriTetra12 Civ
PtriTetra2 Civ
PtriTetra4 Civ
PtriTetra5 Civ
PtriTetra9 Civ
PtriTricu1 Civ
PtriTricu2 Civ
PtriT3168BM Gui
PtriDag41 Civ
PtriInc20 Civ
PtriInc6 Civ
Phataginus tricuspis Cluster 1
PtetTetra10 Civ
PtetTetra7 Civ
PtetTetra8 Civ
PtetCAM085 Cam
PtetT612 Cam
PtetT614 Cam
PtetT616 Cam
PtetYAO001 Cam
Phataginus tetradactyla Cluster 1
SgigT2972 DRC
SgigT2973 DRC
SgigCAM011 Cam
SgigT611 Cam
SgigT613 Cam
SgigT615 Cam
Smutsia gigantea Cluster 1
0.01

Figure_S19. Optimal 12S Neighbour Joining tree of Pholidota bushmeat samples.

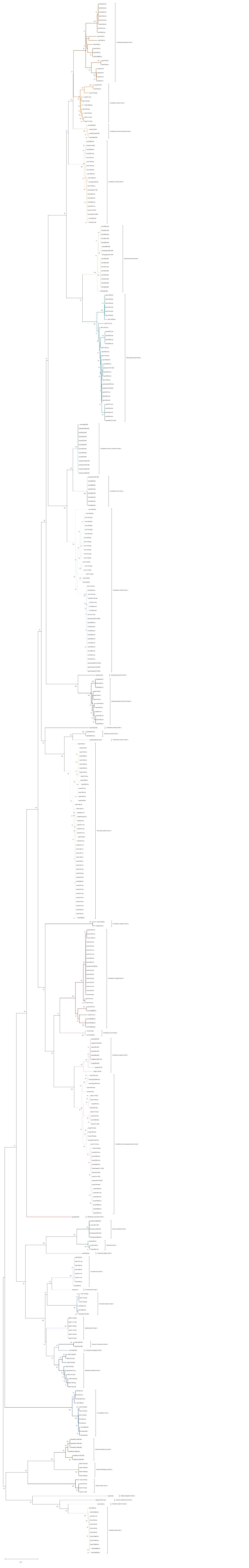

Figure_S20. Optimal COI Neighbour Joining tree of Primates bushmeat samples.

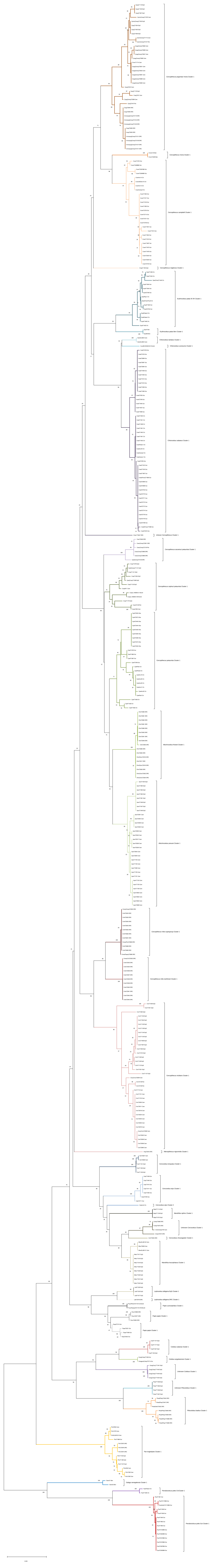

Cercopithecus pogonias/ mona Cluster 1
Cercopithecus mona Cluster 1
Cercopithecus campbelli Cluster 1
Cercopithecus neglectus Cluster 1
Erythrocebus patas W Afr Cluster 1
Erythrocebus patas Ben Cluster 1
Chlorocebus tantalus Cluster 1
Chlorocebus cynosuros Cluster 1
Chlorocebus sabaeus Cluster 1
Unkown Cercopithecus Cluster 1
Cercopithecus ascanius/ petaurista Cluster 1
Cercopithecus cephus/ petaurista Cluster 1
Cercopithecus petaurista Cluster 1
Allochrocebus lhoesti Cluster 1
Allochrocebus preussi Cluster 1
Cercopithecus mitis supergroup Cluster 1
Cercopithecus mitis stuhlmani Cluster 1
Cercopithecus nictitans Cluster 1
Allenopithecus nigroviridis Cluster 1
Cercocebus torquatus Cluster 1
Cercocebus atys Cluster 1
Cercocebus atys Cluster 2
Mandrillus sphinx Cluster 1
Unknown Cercocebus Cluster 1
Cercocebus chrysogaster Cluster 1
Mandrillus leucophaeus Cluster 1
Lophocebus albigena EqG Cluster 1
Lophocebus albigena DRC Cluster 1
Papio cynocephalus Cluster 1
Papio papio Cluster 1
Papio papio Cluster 1
Colobus satanas Cluster 1
Colobus angolanensis Cluster 1
Unknown Colobus Cluster 1
Unknown Piliocolobus Cluster 1
Piliocolobus badius Cluster 1
Pan troglodytes Cluster 1
Galago senegalensis Cluster 1
Perodocicticus potto ClvCluster 1
Perodocicticus potto Gui Cluster 1

Figure_S21. Optimal Cytb Neighbour Joining tree of Primates bushmeat samples.

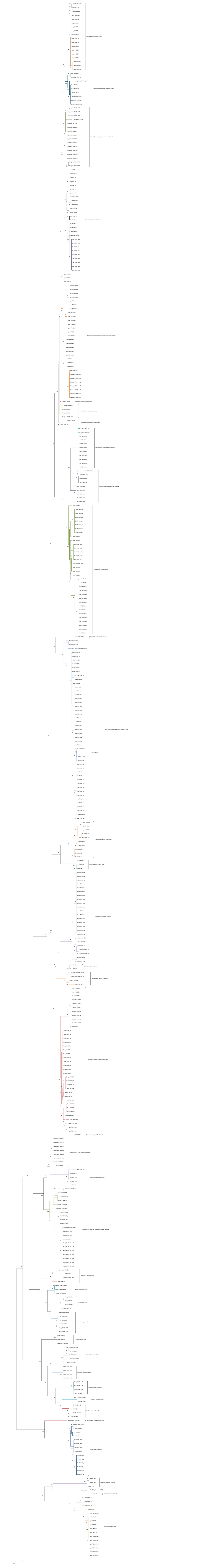

Figure_S22. Optimal 16S Neighbour Joining tree of Primates bushmeat samples.

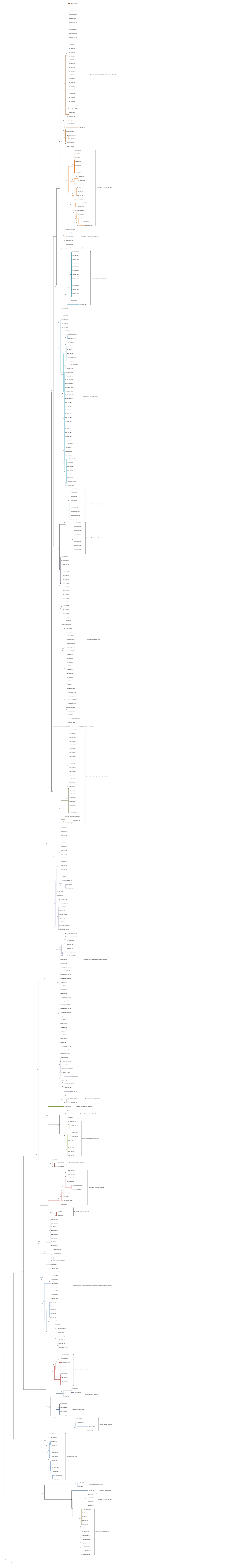

Figure_S23. Optimal 12S Neighbour Joining tree of Primates bushmeat samples.

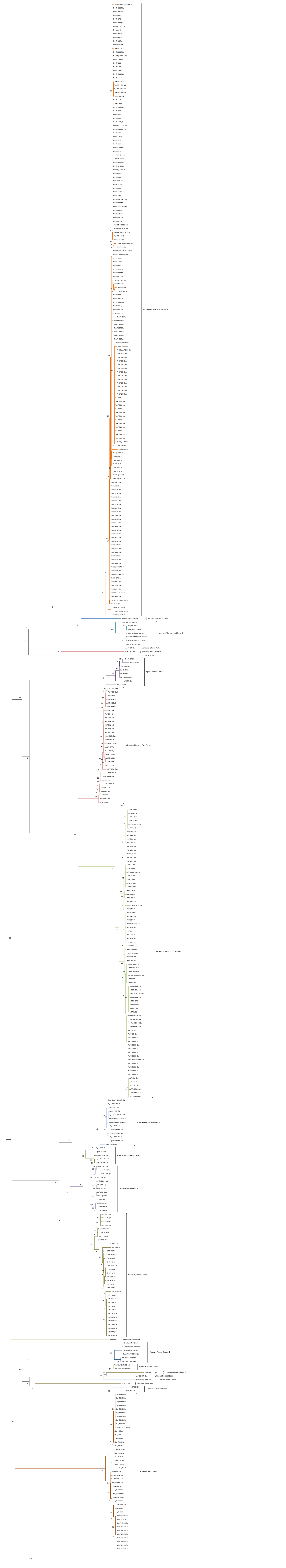

Thryonomys swinderianus Cluster 1
Unknown Thryonomys sp Cluster 1
Unknown Thryonomys Cluster 2
Anomalurus derbianus Cluster 1
Anomalurus beecrofti Cluster 1
Hystrix cristata Cluster 1
Atherurus africanus LC Afr Cluster 1
Atherurus africanus W Afr Cluster 1
Unknown Cricetomys Cluster 1
Cricetomys gambianus Cluster 1
Cricetomys sp3 Cluster 1
Cricetomys sp1 Cluster 1
Arvicanthis rufinus Cluster 1
Unknown Rodent CLuster 1
Unknown Rodent Cluster 2
Unknown Rodent Cluster 3
Unknown Rodent CLuster 4
Unknown Rodent Cluster 5
Unknown Sciuridae Cluster 1
Heliosciurus rufobrachium Cluster 1
Xerus erythropus Cluster 1
0.050

Figure_S24. Optimal COI Neighbour Joining tree of Primates bushmeat samples.

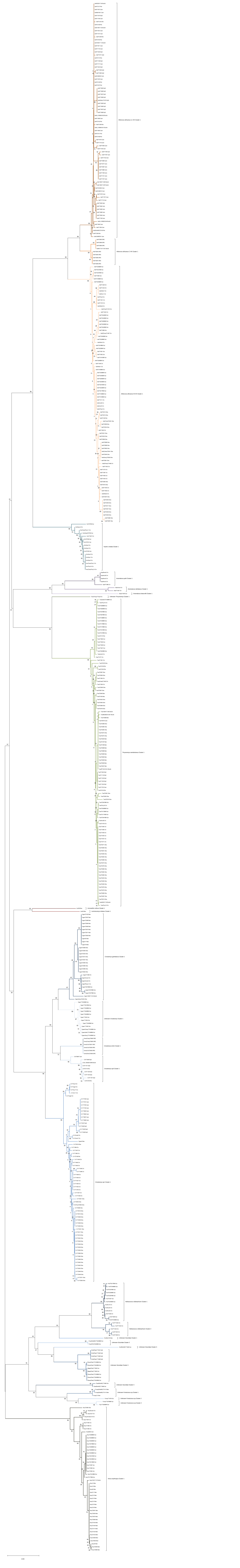

Atherurus africanus LC Afr Cluster 1
Atherurus africanus C Afr Cluster 1
Atherurus africanus W Afr Cluster 1
Hystrix cristata Cluster 1
Anomalurus pelii Cluster 1
Anomalurus derbianus Cluster 1
Anomalurus beecrofti Cluster 1
Unknown Thryonomys Cluster 1
Thryonomys swinderianus Cluster 1
Arvicanthis rufinus Cluster 1
Lemniscomys striatus Cluster 1
Cricetomys gambianus Cluster 1
Unknown Cricetomys Cluster 1
Cricetomys emini Cluster 1
Cricetomys sp3 Cluster 1
Cricetomys sp1 Cluster 1
Heliosciurus rufobrachium Cluster 1
Heliosciurus rufobrachium Cluster 2
Unknown Sciuridae Cluster 1
Unknown Sciuridae Cluster 2
Unknown Sciuridae Cluster 3
Unknown Sciuridae Cluster 3
Unknown Sciuridae Cluster 4
Unknown Funisciurus sp Cluster 1
Unknown Funisciurus sp Cluster 2
Unknown Funisciurus sp Cluster 2
Xerus erythropus Cluster 1

Figure_S25. Optimal Cytb Neighbour Joining tree of Primates bushmeat samples.

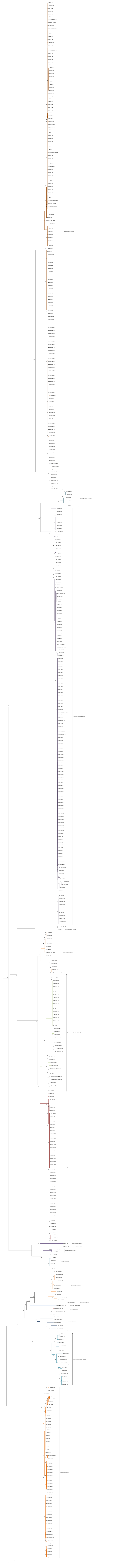

Figure_S26. Optimal 16S Neighbour Joining tree of Primates bushmeat samples.

Figure_S27. Optimal 12S Neighbour Joining tree of Primates bushmeat samples.

Figure S28

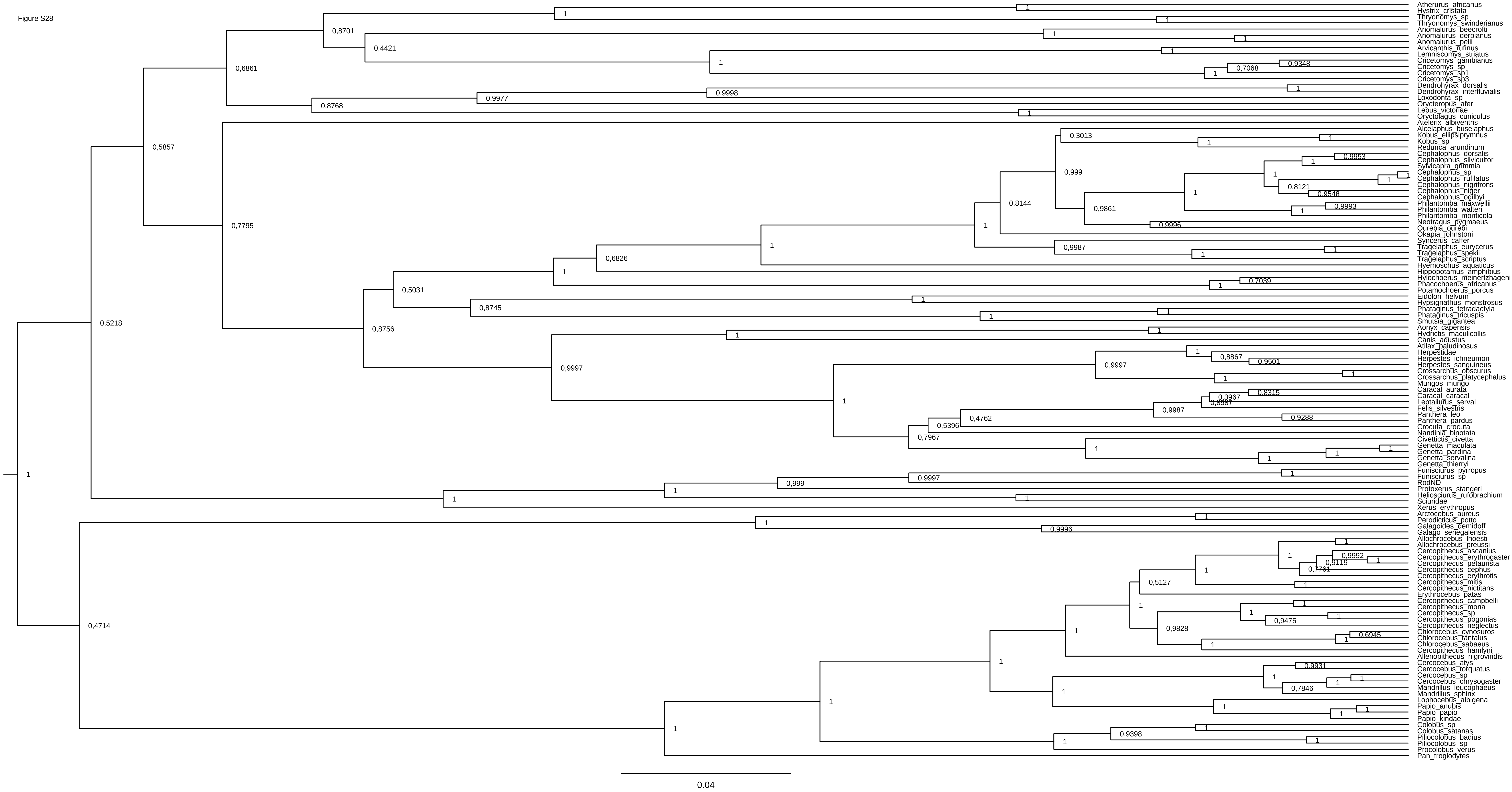
Atherurus_africanus
Hystrix_cristata
Thryonomys_sp
Thryonomys_swinderianus
Anomalurus_beecrofti
Anomalurus_derbianus
Anomalurus_pelii
Arvicanthis_rufinus
Lemniscomys_striatus
Cricetomys_gambianus
Cricetomys_sp
Cricetomys_sp1
Cricetomys_sp3
Dendrohyrax_dorsalis
Dendrohyrax_interfluvialis
Loxodonta_sp
Orycteropus_afer
Lepus_victoriae
Oryctolagus_cuniculus
Atelerix_albiventris
Alcelaphus_buselaphus
Kobus_ellipsiprymnus
Kobus_sp
Redunca_arundinum
Cephalophus_dorsalis
Cephalophus_silvicultor
Sylvicapra_grimmia
Cephalophus_sp
Cephalophus_rufilatus
Cephalophus_nigrifrons
Cephalophus_niger
Cephalophus_ogilbyi
Philantomba_maxwellii
Philantomba_walteri
Philantomba_monticola
Neotragus_pygmaeus
Ourebia_ourebi
Okapia_johnstoni
Syncerus_caffer
Tragelaphus_eurycerus
Tragelaphus_spekii
Tragelaphus_scriptus
Hyemoschus_aquaticus
Hippopotamus_amphibius
Hylochoerus_meinertzhageni
Phacochoerus_africanus
Potamochoerus_porcus
Eidolon_helvum
Hypsignathus_monstrosus
Phataginus_tetradactyla
Phataginus_tricuspis
Smutsia_gigantea
Aonyx_capensis
Hydrictis_maculicollis
Canis_adustus
Atilax_paludinosus
Herpestidae
Herpestes_ichneumon
Herpestes_sanguineus
Crossarchus_obscurus
Crossarchus_platycephalus
Mungos_mungo
Caracal_aurata
Caracal_caracal
Leptailurus_serval
Felis_silvestris
Panthera_leo
Panthera_pardus
Crocuta_crocuta
Nandinia_binotata
Civettictis_civetta
Genetta_maculata
Genetta_pardina
Genetta_servalina
Genetta_thierryi
Funisciurus_pyrropus
Funisciurus_sp
RodND
Protoxerus_stangeri
Heliosciurus_rufobrachium
Sciuridae
Xerus_erythropus
Arctocebus_aureus
Perodicticus_potto
Galagoides_demidoff
Galago_senegalensis
Allochrocebus_lhoesti
Allochrocebus_preussi
Cercopithecus_ascanius
Cercopithecus_erythrogaster
Cercopithecus_petaurista
Cercopithecus_cephus
Cercopithecus_erythrotis
Cercopithecus_mitis
Cercopithecus_nictitans
Erythrocebus_patas
Cercopithecus_campbelli
Cercopithecus_mona
Cercopithecus_sp
Cercopithecus_pogonias
Cercopithecus_neglectus
Chlorocebus_cynosuros
Chlorocebus_tantalus
Chlorocebus_sabaeus
Cercopithecus_hamlyni
Allenopithecus_nigroviridis
Cercocebus_atys
Cercocebus_torquatus
Cercocebus_sp
Cercocebus_chrysogaster
Mandrillus_leucophaeus
Mandrillus_sphinx
Lophocebus_albigena
Papio_anubis
Papio_papio
Papio_kindae
Colobus_sp
Colobus_satanas
Piliocolobus_badius
Piliocolobus_sp
Procolobus_verus
Pan_troglodytes
0.04
